# An integrated Bayesian framework for multi-omics prediction and classification

**DOI:** 10.1101/2022.11.06.514786

**Authors:** Himel Mallick, Anupreet Porwal, Satabdi Saha, Piyali Basak, Vladimir Svetnik, Erina Paul

## Abstract

With the growing commonality of multi-omics datasets, there is now increasing evidence that integrated omics profiles lead to the more efficient discovery of clinically actionable biomarkers that enable better disease outcome prediction and patient stratification. Several methods exist to perform host phenotype prediction from crosssectional, single-omics data modalities but decentralized frameworks that jointly analyze multiple time-dependent omics data to highlight the integrative and dynamic impact of repeatedly measured biomarkers are currently limited. In this article, we propose a novel Bayesian ensemble method to consolidate prediction by combining information across several longitudinal and cross-sectional omics data layers. Unlike existing frequentist paradigms, our approach enables uncertainty quantification in prediction as well as interval estimation for a variety of quantities of interest based on posterior summaries. We apply our method to four published multi-omics datasets and demonstrate that it recapitulates known biology in addition to providing novel insights while also outperforming existing methods in estimation, prediction, and uncertainty quantification. Our open-source software is publicly available at https://github.com/himelmallick/IntegratedLearner.

## 1 INTRODUCTION

Multi-omics is an exciting and emerging field in biomedical sciences that has taken center stage in recent years, facilitating systems biology investigation at an unprecedented scale ^1^. These ever-increasing omics measurements assessed simultaneously in the same biological specimens (e.g., bulk and single-cell transcriptomics, metagenomics and metatranscriptomics, metabolomics and proteomics, and spatial omics and imaging, among others) have led to an explosion of high-throughput multi-modal data, allowing researchers to pinpoint relevant biomarkers across multiple biological entities ^2^. Traditional single-layer analyses of multi-omics data are doomed to be less powerful than integrative approaches and the longstanding interest in addressing this information gap has, therefore, led to a fair amount of recent work aiming to integrate multiple omics modalities for deriving better insights into the underlying biological mechanisms.

Many challenges exist with respect to integrating multiple omics data layers and translating the information to impact human health. For example, many standard single-omics analysis methods from the literature cannot be directly combined to achieve full mechanistic insight without falling prey to false positive or false negative results. This is particularly because different omics modalities have inherent data differences that need to be taken into account during integration and inference. This is further exacerbated by the technical nature of the associated data, which are typically noisy, heterogeneous, and high-dimensional with confounding effects unique to each individual layer (e.g., platform-specific batch effects). A gap thus remains to efficiently combine related features in multi-omics data integration while maintaining sensitivity and controlling spurious association reporting - one of the key hurdles in accurate translational applications of integrative omics to human health.

Despite being a well-studied problem supported by a fair amount of unsupervised algorithms ^3^, multi-omics data integration remains one of the most computationally and biologically challenging problems in the supervised setting, wherein one attempts to discover clinically actionable multi-omics biomarkers predictive of a disease or health outcome. Our use case in this study thus goes far beyond unsupervised approaches in that in addition to combining various key layers, our end goal is also to achieve a holistic understanding of how multiple omics data modalities across biological domains explain or characterize a known phenotype or clinical outcome, which can potentially form the statistical basis to stratify patients for therapeutic interventions in addition to streamlining the discovery of new biomarkers.

Recently, several methods have been proposed for supervised multi-omics data integration. Ghaemi *et al.* (2019) ^4^ used an elastic net stacked generalization approach to integrating multiple omics modalities towards predicting a continuous outcome. Stelzer *et al.* ^5^ similarly considered a combination of LASSO and fused LASSO regression to predict a continuous outcome using integrated trajectories of multiple omics data layers. Franzosa *et al.* (2019) ^6^ used a concatenated random forest approach to predict a categorical outcome using pooled multi-omics profiles. Singh *et al.* (2019) ^3^ proposed a sparse generalized canonical correlation analysis approach to classify subjects and identify variables based on a lower-dimensional representation of the multiomics measurements. More recent advances have focused on deep learning methods for generating highly accurate predictions from multi-omics features ^7,8^, yet they typically rely on non-interpretable ‘black box’ algorithms infeasible for most translational applications.

Among the pros and cons of these individual frequentist methods, the stacked generalization approach, as considered by several authors ^4,9,5^, lacks a proper cross-validation step during the meta-learning stage, potentially introducing overfitting and data leakage during training ^10^. Singh *et al.* (2019) ^3^‘s approach, on the other hand, can only be applied to a categorical outcome and the associated implementation (DIABLO) requires non-calibrated parameter specification and expensive cross-validation, which induces impractically high computation burden to achieve realistic prediction performance. In addition, DIABLO relies on a non-probabilistic majority voting to predict class labels, leading to a biased interpretation of the final prediction performance. None of these frameworks formally provide an approach to inference, and, they are not flexible in quantifying uncertainties in prediction and estimation. This means that practitioners and machine learners typically must deploy the resulting prediction and classification algorithms without a quantification of its uncertainty or without the capability to call out actionable features for basic biological applications.

We address both these issues by providing a flexible, fully Bayesian ensemble approach that combines prediction across multiple omics layers while simultaneously accounting for the nuances of the individual data modalities. We have implemented this method as *IntegratedLearner*; it uses a Bayesian additive regression trees approach ^11^ at each individual layer and uses non-negative least squares optimization or rank loss minimization ^12^ to optimally combine predictions across layers. In this study, we also use a two-stage joint modeling approach to seamlessly integrate longitudinal multi-omics biomarkers in the prediction algorithm. To the best of our knowledge, the two-stage Bayesian ensemble approach presented here is the first to consider such aspects of longitudinal multi-omics biomarkers by jointly analyzing all time points across data types. In addition to reporting interpretable predictive features that are (potentially) non-linearly related to the outcome, the proposed Bayesian approach provides both point and interval estimation for a variety of quantities of interest as an automatic byproduct of the corresponding Markov Chain Monte Carlo (MCMC) procedure. We demonstrate the functionalities of our proposed framework by applying it to four publicly available multi-omics datasets to identify relevant biomarkers of various disease and health outcomes and conduct simulation experiments to further validate the method in a range of realistically complex scenarios.

The remainder of this article is organized as follows. In Section 2, we describe the *IntegratedLearner* methodology. Next, we present our numerical results in Section 3, where we also compare the performance of our methodology to the performance of other published methods. Finally, we conclude with a brief discussion of our results in Section 4 along with potential next steps in this area of research.

## 2 METHODS

We first describe the *IntegratedLearner* algorithm in Section 2.1. We then introduce the prior formulation along with the algorithms for estimating the posterior summaries based on Bayesian additive regression trees in Section 2.2. We describe the associated data integration strategy to combine predictions from longitudinal and cross-sectional multi-omics biomarkers in Section 2.3. We also describe a simple yet effective and interpretable approach to combine predictions (Section 2.4) and conclude by summarizing the major contributions of this article (Section 2.5).

### 2.1 The *IntegratedLearner* Algorithm

*IntegratedLearner* identifies predictive biomarkers from potentially heterogeneous multiple omics datasets co-indexed along one axis. This co-indexing is referred to here as the “samples” axis, whereas, we use “features” and “profiles” to refer to the quantification of multi-omics measurements from high-throughput experiments (e.g., gene expression counts, microbial taxonomic and functional profiles, and protein or metabolite abundances, among others). For a set of omics datasets containing measurements that describe the same set of samples, the *IntegratedLearner* algorithm proceeds by 1) correcting for potential confounding variables and batch effects per data layer (if desired), 2) applying pre-processing and independent filtering to each dataset to generate quality-controlled feature sets (if specified), 3) fitting a per-layer Bayesian additive regression trees method ^11^ to predict outcome (base-learners), and 4) combining the layer-wise out-of-sample predictions using non-negative least squares or rank loss minimization (meta-learner) to generate final predictions based on all available data points (**Fig. 1**). The final model, essentially, is a weighted average of the individual tree models where the weights are derived from the per-layer Bayesian non-parametric ensembles.

**FIGURE 1.**
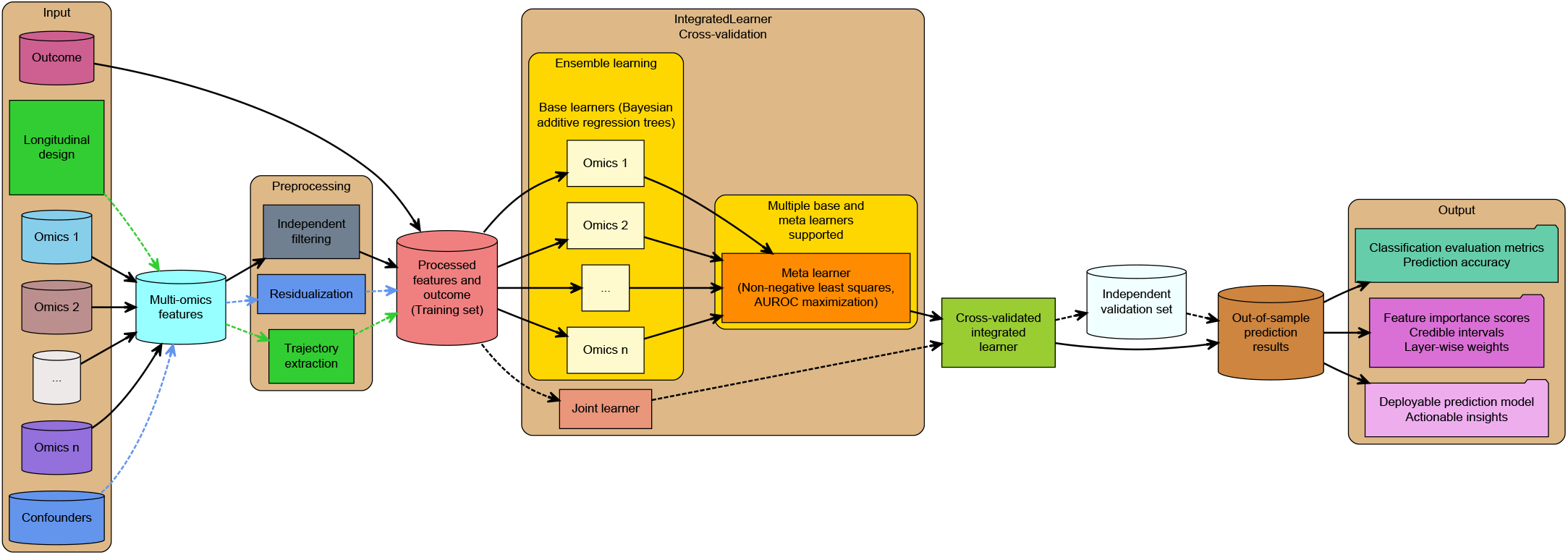
*IntegratedLearner* is an integrated Bayesian statistical framework for end-to-end biomarker discovery from cross-sectional and longitudinal multi-omics data. In order to facilitate multi-omics prediction and classification of a clinical outcome (continuous or categorical), it uses a Bayesian ensemble approach to combine predictions across multiple omics layers. For each omics modality, beginning with individually quality-controlled training data, it identifies a minimal set of features most predictive of the outcome based on rigorously cross-validated Bayesian additive regression trees models. These individual learners are then combined to build the optimally weighted combination of predictions, where optimality is defined by a user-specified objective function, such as minimizing the mean squared error or maximizing the area under the receiver operating characteristic curve. In the presence of repeated measures and covariates, *IntegratedLearner* optionally extracts random effects and covariate-adjusted residuals prior to ensemble learning. If requested, it further provides predictions based on a concatenated model to facilitate comparisons with the default ensemble approach. Upon training and validation, out-of-sample performance measures are summarized and feature importance scores as well as posterior summaries (for calculating credible and prediction intervals) are returned for further downstream analysis. In addition to the default base- and meta-learners, multiple machine learning algorithms are supported (**Fig. S1**). Dotted and solid arrows respectively represent optional and default and/or recommended steps in the workflow.

The base-learner fitting process optimizes a small number of features per layer and the best model for a particular layer is selected based on the lowest cross-validated error obtained via a rigorous cross-validation procedure. In the meta-learning stage, out-of-sample predictions from the first stage are used as features to estimate the layer weights. In the presence of longitudinal biomarkers, our method uses a two-stage joint modeling approach (**Section 2.3**), which consists of a per-feature linear mixed effects model for the longitudinal data, wherein the predicted values of the random intercepts are used as “features” in the *IntegratedLearner* algorithm. Upon training and validation, out-of-sample performance measures are summarized and feature importance scores as well as posterior summaries (for calculating credible and prediction intervals) are returned for further downstream analysis.

### 2.2 Bayesian Additive regression trees

The building block of the *IntegratedLearner* algorithm is a per-layer Bayesian additive regression trees (BART) method ^11^ which is described in detail in Tan *et al.* (2019) ^13^. Briefly, BART is a nonparametric tree ensemble method that robustly estimates a regression function especially when nonlinear relationships and/or interactions are present ^11^. BART is a Bayesian “sum-of-trees” model in which each of the individual trees explains only a small portion of the outcome variability and each tree is constrained by a regularization prior to be a weak learner. The trees work like small cogs in a machine with assigned roles, but work as an ensemble to perform more complex operations ^11^. These individual trees may have a larger bias than a single complex tree, but they collectively estimate the unknown regression function with a lower bias than the individual weak learners. Fitting and inference are accomplished by means of an iterative backfitting Markov Chain Monte Carlo (MCMC) algorithm that generates samples from a posterior. BART enables full posterior inference including point and interval estimates of the unknown regression function as well as the variable importance measures of individual predictors.

In this subsection, we briefly review the BART method which is a non-parametric Bayesian method that uses a Bayesian ‘sum trees’ model that enables full posterior inference including point and interval estimates of the unknown regression function. Inspired by ensemble methods, each tree in this model is constrained by a regularization prior allowing it to be a weak learner. Fitting and inference are then accomplished via an iterative Bayesian back-fitting MCMC algorithm that generates samples from the posterior. The successive post-burn-in iterations of the back-fitting MCMC algorithm thus form a full draw of MCMC samples generated from the posterior distribution over the ‘sum of trees’ model space. The posterior mean estimate of any input feature is simply obtained by averaging over the consecutive ‘sum of trees’ model draws. Feature selection is made possible by calculating the relative frequencies of the corresponding components appearing in the ‘sum of trees’ model iterations. The corresponding relative frequencies help in explaining the relative importance of the input features in determining the variation in the outcome.

To explain the form of a ‘sum of trees’ model we start with a single tree model. Let *z*_*i*_ denote a continuous outcome and *x*_*i*_ denote the vector of covariates modeled by the structure *z*_*i*_ = *g*(*x*_*i*_; *T, M*) +ϵ_*i*_. Notationally, T denotes a binary tree consisting of interior and terminal nodes. A branch decision rule at each interior node typically splits the predictor space into two regions {*x* ∈ *S*} vs {*x* ∉*S*}, where S is the covariate space. Let *M* = {*μ*_1_, *μ*_2_,, *μ*_*b*_} denote a set of functional values associated with the *b* terminal nodes. For a given T and M, we use *f* (*x*_*i*_; *T, M*) to denote a function that assigns a *μ*_*i*_ E *M* to *x*_*i*_. Thus,

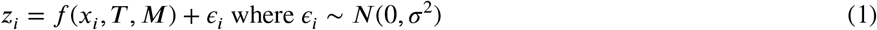

is a single tree model and the BART model for continuous outcome *z*_*i*_ can be expressed as

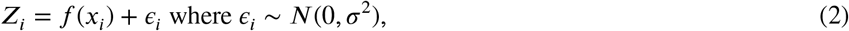

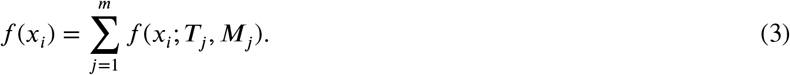

The BART model for the classification of a categorical outcome *y*_*i*_ can be reformulated from the continuous model using a probit transformation as follows:

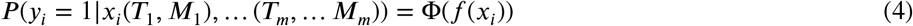

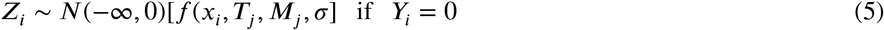

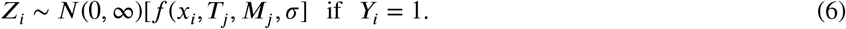

For the Bayesian specification of the ‘sum of trees’ model, we specify regularization priors over the parameters of the BART model, namely, (*T*_1_, *M*_1_), …, (*T*_*m*_, *M*_*m*_) and σ. Priors are carefully chosen to regularize the fit and to curtail strong individual tree effects, specified as follows:

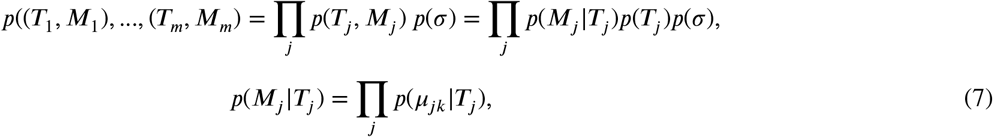

where *μ*_*j k*_ ∈*M*_*j*_. Following Chipman *et al.* (2010) ^11^, we specify *p*(*T*_*j*_) to represent three aspects, (i) the probability that a node at depth *d* is non-terminal is calculated as *α*(1 + *d*)^− ζ (^ where *α* ∈ (0, 1) and ζ ∈ [0, ∞), (ii) the choice of the splitting covariate at each interior node is uniformly distributed over the set of available covariates, and (iii) the choice of a branching rule given a covariate at the interior node also follows a uniform distribution over the discrete set of available splitting values. We consider the default values recommended by Chipman *et al.* (2010) ^11^: *α* = 0.95, *ζ*= 2. For *p*(*M*_*j*_ \*T*_*j*_) = ∏_*j*_*p*(*μ*_*jk*_|*T*_*j*_), we use the conjugate normal prior *μ*_*jk*_ ∼ *N*(0, 2.25/*m*) on the values of the terminal nodes. This prior has the effect of curbing strong individual effects of the trees so that every tree forms only a small part of the entire ‘sum of trees’ model, thus optimally eliminating the chance of overfitting at individual data layers. For the probit transformation, the same algorithm proceeds after setting σ to 1.

### 2.3 Two-stage model for longitudinal multi-omics biomarkers

In the presence of longitudinal or clustered multi-omics data, we adhere to a two-stage analysis framework introduced by Horrocks *et al.* (2009) ^14^ for binary outcome prediction and extend it to the more general case of multi-omics prediction and classification. Let *X* _*i*_= (*X*_*i*1_, X_*i*2_, …, X_*ip*_)’s represent the longitudinal multi-omics biomarkers for individual *i, i* = 1, 2, … , *n*, at measurement times *t*_*i*_ = (*t*_*i*1_, … , *t*_*ip*_)′. Before calling the individual base-learners, we fit a per-feature linear mixed effects model following layer-specific normalization and covariate adjustment as:

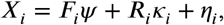

where *F*_*i*_ is the design matrix corresponding to the vector of unknown fixed-effects parameters (*Ψ*) representing the per-layer confounders, *R*_*i*_ is the design matrix corresponding to the random coefficients κ_*i*_, and *η*_*i*_ is the vector of random noise for individual *i*. Of note, in this manuscript, we consider the simple case of random intercepts as random coefficients (κ_*i*_) but the methodology is applicable to a more general case where the random design matrix *R*_*i*_ may contain random effects terms in the form of splines or other functional curves. Assuming independent multivariate normality of the random coefficients, we estimate the random effects *κ*_*i*_ which are used as second-stage features in the *IntegratedLearner* framework (**Fig. 1**) to generate subject-level predictions of the outcome of the interest. The embedding of this strategy in the paradigm of linear mixed models enables the treatment of diverse data types in a unified setting while also allowing the extraction of subject-level longitudinal trajectories in the form of random coefficients, not captured by existing cross-sectional machine learners.

### 2.4 Simple and interpretable meta-model

In the meta-learning stage, all out-of-sample predictions from the first stage are used as features. Assuming a *J* - fold crossvalidation (CV) procedure during the first-stage learning for e ach o f t he *K* m odalities, we a ssemble t he o ut-of-sample CV predictions to get ‘Level 1’ data:

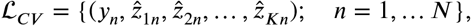

where 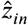 is the predicted values (for continuous outcomes) or the predicted probabilities (for binary outcomes) from the *k*^*th*^ modality for the *n*^*th*^ data point. For continuous outcomes, we use a non-negative least squares optimization to learn the contribution of each layer to the response **Y** leading to the optimal weights (*α*_*j*_ ‘s) for combining predictions, i.e.,

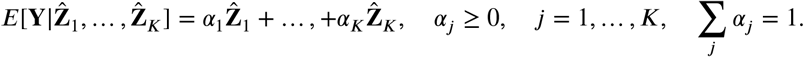

For the binary response, we similarly model 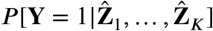 as

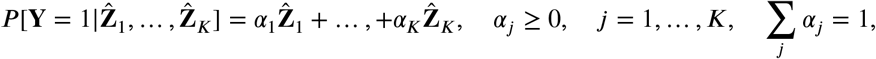

and minimize rank loss defined by (1 − *AUC*), leading to the following objective function:

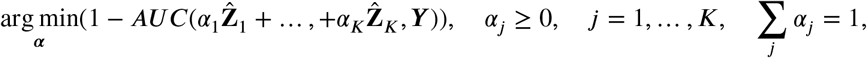

where AUC is the area under the receiver operating characteristic (ROC) curve. The above procedure thus allows us to estimate an optimal combination of layer-specific predictions in the meta-learning stage. The estimated weights can be interpreted as the relative contributions of individual modalities in the final prediction, which can potentially serve as a cost-effectiveness indicator for future multi-omics data collection efforts. In contrast to existing stacked generalization method ^4,9,5^, where additional crossvalidation is typically used during the second-stage learning (ensuring that the same set of data points are held out at all levels of the analysis), we bypass a second cross-validation step and instead re-fit the individual base-learners on the entire samples and combine these predictions with the estimated cross-validated weights to avoid potential overfitting and data leakage during training ^10,15^.

To calculate the weighted average posterior samples, we use the estimated weights from the meta-model and the associated layer-wise posterior samples from the first-stage learning, which facilitates the estimation of a variety of downstream quantities such as feature importance scores, credible intervals, and prediction intervals, among others. *IntegratedLearner* thus provides a systematic procedure to quantify uncertainty in predictions; by constructing individual base-learners during the first-stage learning, *IntegratedLearner* avoids dealing with different scales, collection bias, and noise in different data types, whereas, by integrating data in a non-linear fashion, *IntegratedLearner* takes advantage of the common as well as complementary information in different data types by efficiently fusing these into one simple and interpretable meta-learner that represents the full spectrum of the underlying data.

### 2.5 Summary of major contributions

Despite its wide use in practice, BART has been underutilized in the multi-omics data integration literature ^13^. To the best of our knowledge, the proposed approach is one of the first non-trivial applications of BART in a cross-dataset learning task, especially in the context of integrated analysis of multi-omics data. It is to be noted that a routine application of BART to a concatenated data frame (i.e., simply merging all available datasets into a single matrix for joint modeling) can yield severely biased predictions due to the presence of strong platform-specific confounding factors and heterogeneous batch effects introduced by the differential sizes, modularities, and normalizations of the included datasets ^4^. Further, it is not possible to use BART without non-trivial modifications to handle the increasingly common longitudinal nature of modern multi-omics studies, often entangled with multiple covariates and a mixture of cross-sectional and longitudinal data components across layers ^16^. Finally, the Bayesian model permits effect heterogeneity (across layers) to be regularized separately in each individual layer, making it possible to informatively shrink to homogeneity in the final stage of the analysis. Our work thus represents one of the first attempts to comprehensively combine cross-sectional and longitudinal multi-omics biomarkers, superseding previous efforts by including multiple covariates and longitudinal designs while facilitating seamless uncertainty quantification and interpretable variable selection. The implementation of *IntegratedLearner* is publicly available with source code, documentation, tutorial data, and as an R/Bioconductor software implementation.

## 3 RESULTS

### 3.1 *IntegratedLearner* vastly outperforms existing multi-omics prediction methods

We first assessed the degree to which *IntegratedLearner*’s two-layer model (**Fig. 1**) benefits compared to published multi-omics prediction results based on real data. To this end, we trained a default *IntegratedLearner* model on the same raw gut microbiome multi-omics dataset used in the Franzosa *et al.* (2019) ^6^ study, which consisted of 9171 quality-controlled features from 2 layers (**Table 1**). In this study, stool samples were collected from a cross-sectional cohort of individuals enrolled in the Prospective Registry in IBD Study at MGH (PRISM) in order to characterize the gut metabolic profile and microbiome composition in Inflammatory Bowel Diseases (IBD). This cohort included 155 subjects: 68 with Crohn’s disease (CD), 53 with ulcerative colitis (UC), jointly grouped as IBD, and 34 non-IBD controls. Each stool sample was subjected to metagenomic sequencing followed by the profiling of microbial community taxonomic composition and functional potential. In addition, each sample was analyzed by four liquid chromatography tandem mass spectrometry (LC-MS) methods measuring polar metabolites, lipids, free fatty acids, and bile acids, respectively. In addition to carrying out a holistic investigation of the microbiome–metabolome interface, one of the primary objectives of this study was 1) to assess the power of the metabolomic and microbial layers in classifying IBD status and 2) to investigate whether most IBD trends observed in the discovery cohort replicate in an independent validation cohort (**Table 1**).

**TABLE 1.**
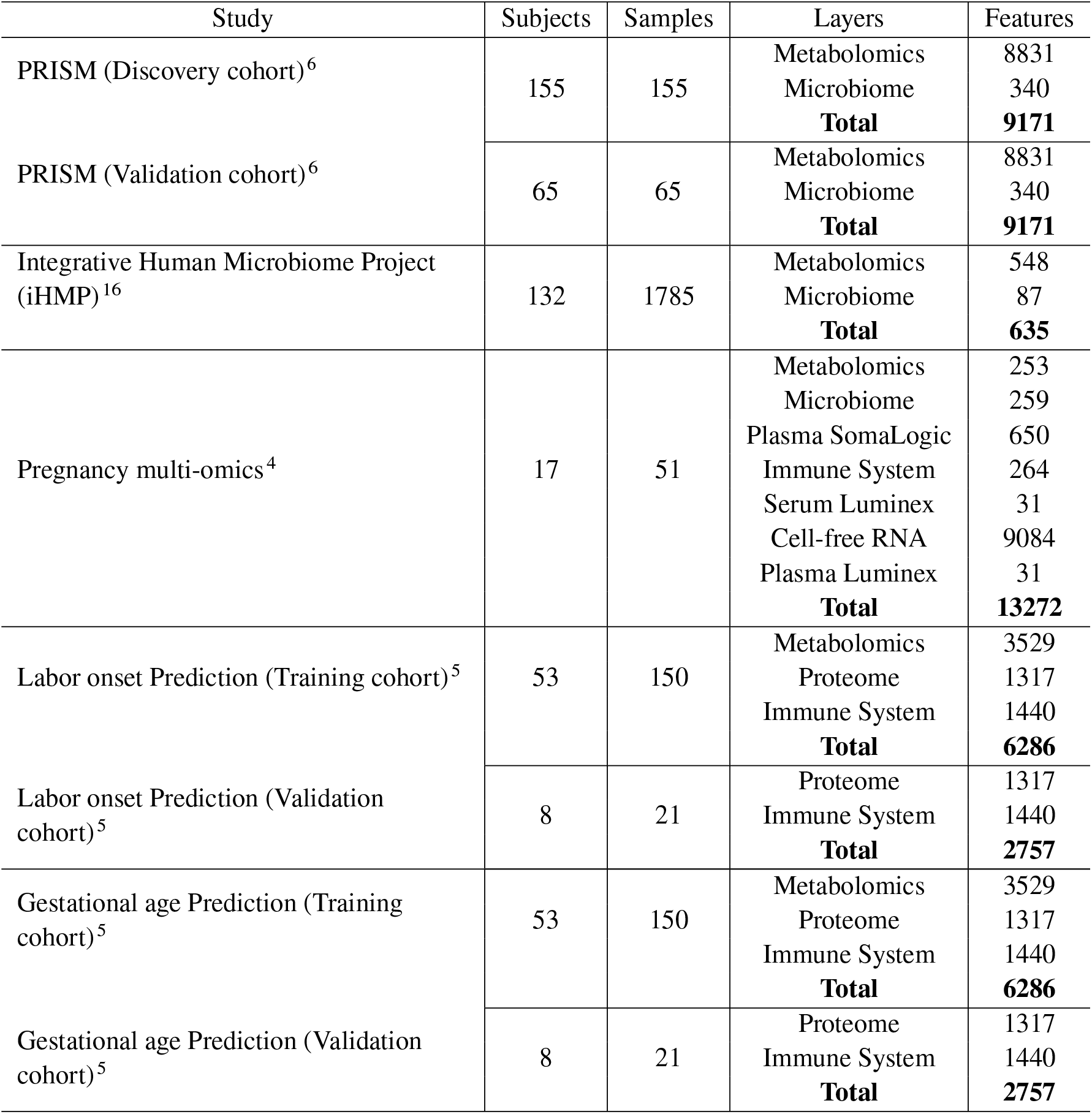
Summary of public multi-omics datasets evaluated in this study.

Franzosa *et al.* (2019) ^6^ observed that both microbial features and metabolites tend to be mutually differentially abundant in IBD and they often covary independently of their mutual covariation with disease. In order to de-emphasize associations that are driven purely by mutual association with covariates unrelated to disease, we trained our IBD classifier on the metabolite and species data after residualizing the effect of potential confounders such as age and antibiotic usage. This residualization step ensures interpretable decision rules while avoiding overly optimistic prediction results caused by co-varying features. To evaluate if differences in metabolite or microbial composition could be used to classify patients according to the IBD phenotype, we learned a default *IntegratedLearner* classifier and compared our method with the concatenated random forest approach from the original study ^6^. Classification performance was evaluated within the PRISM cohort (using 5-fold cross-validation) and between cohorts by training on the entire PRISM cohort and validating on the independent validation cohort.

Several observations are in order. First, as previously observed, the integrated model learned from multi-omics datasets perormed similarly to those using only one type of omics data, which agrees with Franzosa *et al.* ^6^, who also observed that the classifiers trained on metabolite features or microbial species performed similarly to their concatenated counterpart (**Fig. 2**). Second, all the classifiers performed considerably better than random in the task of distinguishing IBD and non-IBD controls (as measured by the AUC, with cross-validation results (AUC 0.93 - 0.96) slightly better than independent validation results (AUC 0.75 - 0.96), indicating that the classifiers generalized well to the validation cohort (**Figs. 2A-B**). Third, while the classifiers trained on metabolite features versus microbial species performed similarly in the discovery cohort, species profiles contributed more to the final prediction despite being considerably smaller in dimension (**Table 1**), suggesting that the single-omics data modularity was not correlated with the corresponding predictive power (**Fig. 2C**), consistent with the literature ^4^. Finally, *IntegratedLearner* significantly outperformed Franzosa *et al.* (2019) ^6^ in external independent validation while also providing valid credible intervals (**Figs. 2D-E**) as well as feature importance intervals (**Fig. S2**) not easily attainable by published methods. We also compared our method with stacked elastic net ^6^ and DIABLO ^3^, which further attests to the superior performance of *IntegratedLearner* across learning tasks (**Table S1**). This finding highlights the benefit of a Bayesian approach and specifically the advantage of using a Bayesian ensemble meta-learning approach as a yardstick for multi-omics classification, as it seems to capture the full range of factors contributing to the outcome or disease status.

**FIGURE 2.**
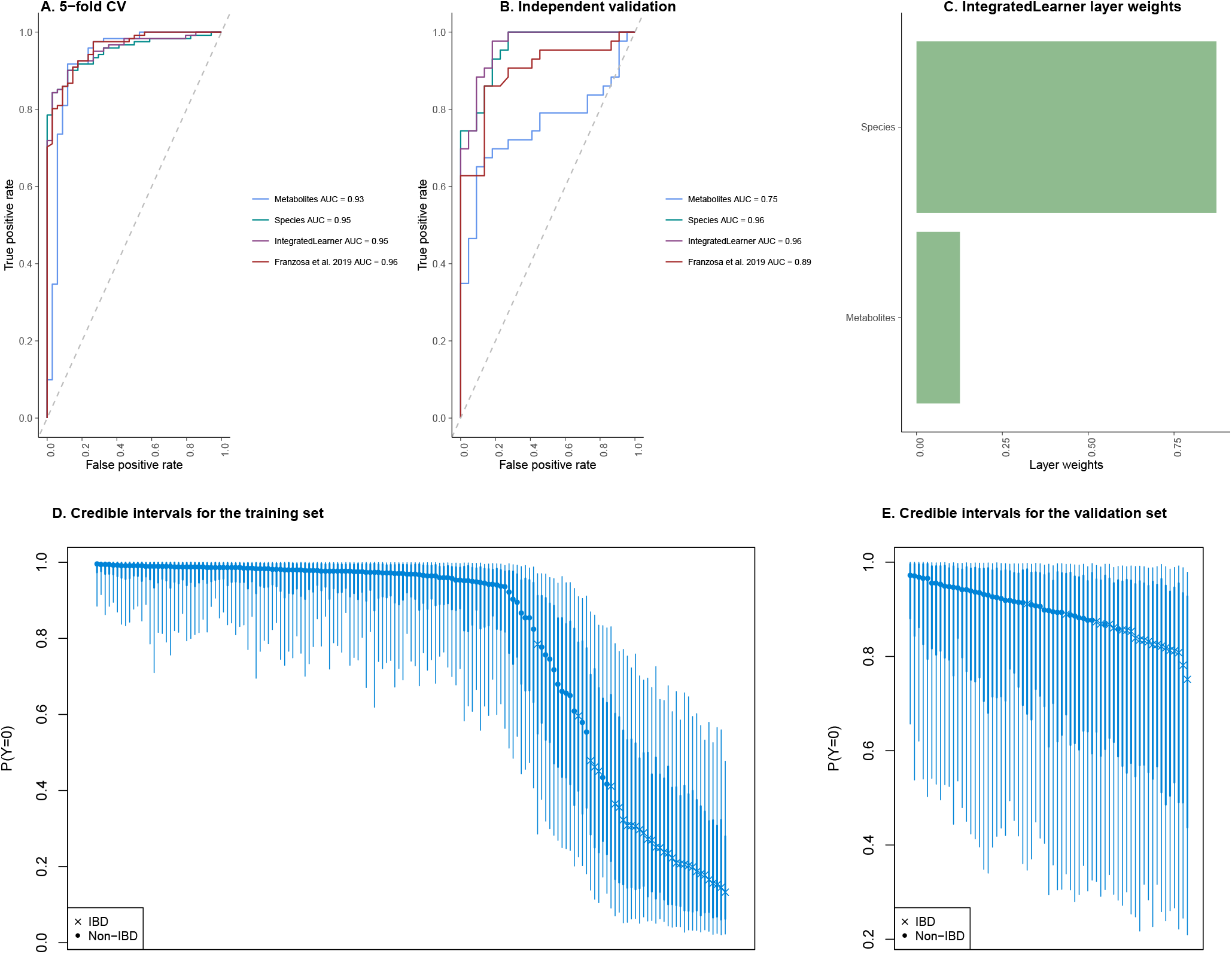
*IntegratedLearner* outperforms state-of-the-art multi-omics prediction methods in classifying IBD patients based on gut microbiome multi-omics features in the PRISM dataset ^6^, as measured by the AUC. Training/testing was carried out within the PRISM training cohort using 5-fold cross-validation (**A**) and external validation on the independent validation cohort (**B**). *IntegratedLearner* also reports layer-specific weights for individual layers (**C**), in addition to the posterior credible intervals for both training (**D**) and test samples (**E**). The thick (thin) line represents 68% (95%) credible intervals; ‘x’ and ‘o’ represent the true class labels and their heights denote the predicted posterior median probabilities of the negative class (i.e., non-IBD).

We hypothesize that the wide 95% credible intervals in the IBD prediction and feature importance (**Figs. 2D-E**; **Fig. S2**) e due to the fact the non-IBD subjects in this population are not perfectly representative of a healthy cohort, thus adding to he significant heterogeneity in an already heterogeneous disease cohort ^6,16,17^. This highlights the importance of quantifying uncertainty in the presence of substantial population heterogeneity as it is misleading to summarize prediction in terms of a single point estimate. Further, we note that there is no clear separation between the predicted posterior median probabilities for IBD vs. non-IBD (**Figs. 2D-E**), which indicates that the best classification threshold in this highly imbalanced problem is possibly far from the default threshold of 0.5 and a follow-up analysis to determine the optimal classification threshold is warranted to better summarize the prediction performance.

### 3.2 *IntegratedLearner* accurately reproduces published multi-omics prediction results

Having demonstrated a clear advantage of the proposed integrative Bayesian approach in a cross-sectional setting, we next examined whether we can recapitulate published multi-omics studies with a longitudinal component. We, therefore, obtained longitudinal multi-omics data from pregnant women in a cohort study at Stanford University ^4^ that aimed to prospectively examine various environmental and biological factors associated with normal and pathological pregnancies. Unlike the PRISM study, this dataset contained repeated measures during pregnancy that allowed assessing important biological adaptations occurring continuously from the early phases of fetal development (first trimester) to the late phases of gestation (third trimester). Considering both the small sample size and the repeated measures aspects of this study, following Ghaemi *et al.* (2019) ^4^, we employed a one-subject-leave-out cross-validation to build prediction models. In order to assess the performance of various machine learning modules available in *IntegratedLearner*, we calculated the coefficient of determination (*R*^2^) based on the observed and cross-validated out-of-sample predicted values of the gestational age.

Ghaemi *et al.* (2019) ^4^ compared several non-Bayesian machine learning methods for predicting gestational age in this dataset and found that their elastic net stack generalization approach achieved the best prediction accuracy in their evaluation. Although we could not validate this claim, possibly due to the differences in the preprocessing steps between the original study and the current study (**Table 1**), our analysis revealed that the proposed Bayesian method remained one of the best-performing methods in accurately predicting gestational age (**Fig. 3**). As before, the integrated models outperformed individual-level base-learners, a phenomenon that was consistent across algorithms.

**FIGURE 3.**
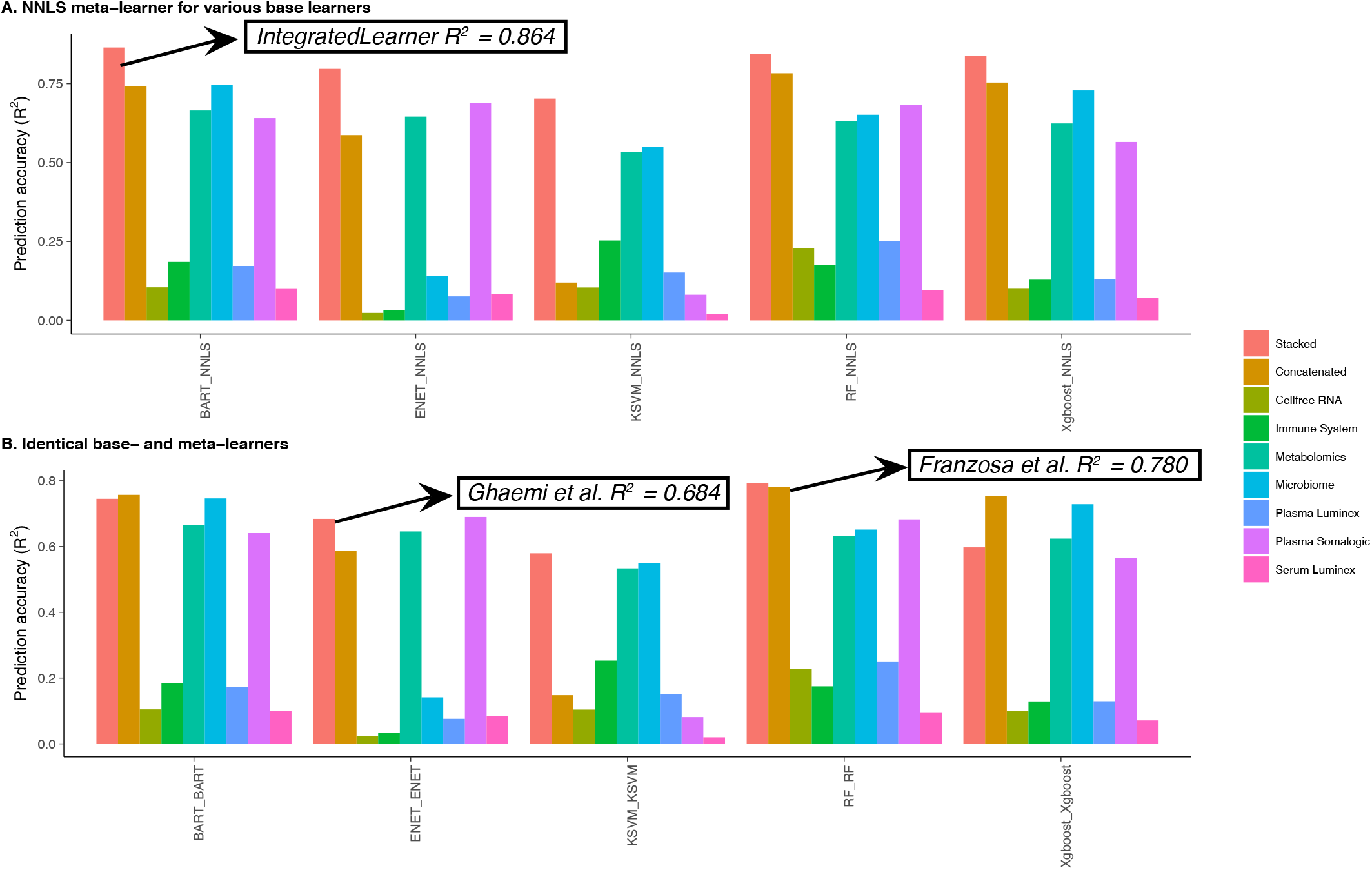
Empirical evaluation of various off-the-shelf and novel machine learning modules implemented in *IntegratedLearner* for the prediction of gestational age in the Ghaemi *et al.* (2019) dataset ^4^. The labels on the x-axis represent various machine learning algorithms used as base- and meta-learner respectively (separated by an underscore). When the meta-learning method is constrained to be non-negative least squares (NNLS) minimization (default parametrization of *IntegratedLearner*, **A**), the proposed method (BART + NNLS) significantly outperforms popular integrated methods while BART performing most favorably in individual layer-wise prediction tasks. **B**. When combined with a computationally demanding non-NNLS metalearner, BART remains one of the best-performing methods, thus highlighting the flexibility of fitting multiple models in a single unified framework.

In agreement with Ghaemi *et al.* (2019) ^4^, we observed that concatenation-based approaches, which are often susceptible to biases introduced by the differential properties of the included datasets, did not exhibit superior performance compared to the ensemble approach. Apart from the integrated level, *IntegratedLearner* maintained similar accuracy at several individual layers including the microbiome, metabolomics, and plasma SomaLogic, which are also the top three most predictive data layers according to the estimated meta-learner weights (**Fig. S3**). Further, in addition to providing easy-to-interpret credible intervals, the Bayesian formulation also provides valid standard errors for the variable importance scores which can be problematic for the frequentist methods (**Fig. S2**). Finally, in addition to the default Bayesian setting, our software implementation *IntegratedLearner* can fit other non-Bayesian base- and meta-learners specified by the users (**Fig. 3**). This option makes *IntegratedLearner* a powerful and viable tool for evaluating a variety of multi-omics-focused machine learning methods in a unified estimation umbrella. Collectively, the results of these validation experiments demonstrate that *IntegratedLearner* can be used to recapitulate findings from real-world multi-omics datasets, while also providing a solid quantification of the various posterior quantities based on geometrically ergodic MCMC samples.

### 3.3 Novel multi-omics biomarker discovery from the integrated Human Microbiome Project

As an illustration of *IntegratedLearner*’s ability to generate novel biological insights from a longitudinal multi-omics study, we consider the problem of predicting the cross-sectional IBD status, in a population of IBD and non-IBD patients, based on longitudinal measurements of metagenomics profiles and metabolomics measurements. A goal of the analysis is to provide, for each subject, an estimated probability of IBD disease. To this end, we applied *IntegratedLearner* to identify relevant IBD-associated microbial features using longitudinal multi-omics data from the Integrative Human Microbiome Project (iHMP) ^16^. The iHMP dataset included 132 individuals recruited in five US medical centers with Crohn’s disease (CD), ulcerative colitis (UC), and non-IBD controls, followed longitudinally for one year with up to 24 time points each (**Table 1**). Integrated multiomics profiling of the resulting 1,785 stool samples generated a variety of data types including metagenome-based taxonomic profiles, metagenomic and metatranscriptomic functional profiles, and biochemical profiles, producing one of the largest publicly available microbial multi-omics datasets to date ^16^. Following the original study, independent filtering was performed before downstream analysis for each of these data modalities.

As mentioned above, unlike the previous case studies, one particularly challenging aspect of this study was the availability of longitudinal multi-omics biomarkers for predicting a cross-sectional outcome (i.e., IBD disease status). To achieve the generality of our approach beyond cross-sectional classification, we first used the iHMP dataset to extract subject-specific random effects using a two-stage joint model approach ^14^ before building the integrated Bayesian machine learner. Specifically, we fit a per-feature linear mixed effects model to the longitudinal data in each layer and extracted the corresponding predicted random intercepts as input for the classifier. To maximize impact, we focused on two layers (microbiome and metabolomics) that had the least amount of missingness across layers while also ensuring the highest number of pairwise complete observations ^16^. Despite the commonality of longitudinal multi-omics studies, to the best of our knowledge, this particular use case has not been described in the multi-omics literature.

Consistent with the previous case studies, we found that the proposed Bayesian method remained one of the best-performing methods outperforming published methods (**Fig. 4A**), with the greatest signal exhibited by the metabolomics layer (**Fig. 4B**). This phenomenon was consistent across algorithms (**Fig. S4**). Similar to the PRISM dataset, we observed wide credible intervals for the IBD classification probabilities (**Fig. S4**). Apart from accurately classifying IBD patients, *IntegratedLearner*’s Bayesian tree model recapitulated several associations from the original study’s univariate approach ^18^. In particular, we observed that a significantly IBD-depleted Roseburia species (*R. hominis*), not captured by the previous analysis, remained one of the most predictive features in our analysis. Notably, “top hits” from both *IntegratedLearner*’s multivariate prediction model and the original study’s univariate approach yielded many overlaps, which broadly manifested as a characteristic increase in facultative anaerobes at the expense of obligate anaerobes, in agreement with the previously observed depletion of butyrate producers in IBD (**Fig. 4C**). In addition to these taxonomic associations, *IntegratedLearner* also identified several biochemical and functional associations such as specific literature-curated metabolites including the enrichment of bile acid-associated products (**Fig. 4C**). These findings suggest that the two-stage approach is able to successfully transfer both inter-individual and within-subject information to generate an accurate and meaningful classification of IBD status. Importantly, this implies that ignoring within-subjects information may bias future attempts to predict a cross-sectional outcome from longitudinal biomarkers, further highlighting that *IntegratedLearner*’s two-stage approach offers a more interpretable and uncertainty-quantified machine learning for future longitudinal multi-omics biomarker studies with a cross-sectional outcome.

**FIGURE 4.**
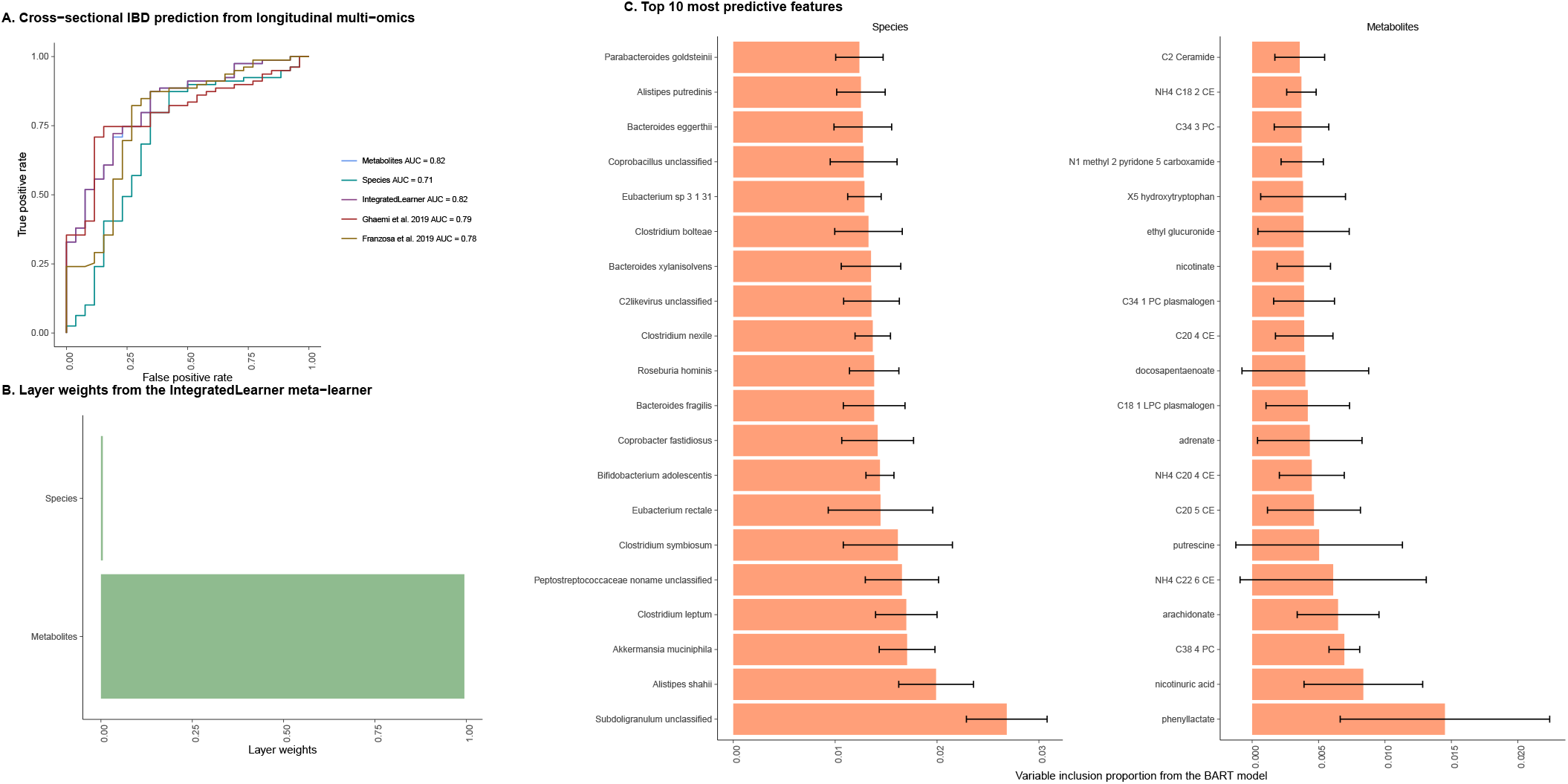
*IntegratedLearner* enables Bayesian joint modelling of longitudinal multi-omics biomarkers for predicting cross-sectional outcomes by undertaking a two-stage strategy that first fits the longitudinal submodel and then plugs in the shared information into the machine learning submodel for outcome prediction. A. We consider the problem of predicting the IBD status (for each subject), in a population of IBD and non-IBD subjects in the integrative Human Microbiome Project (iHMP), based on longitudinal multi-omics measurements ^16^. Our results show that the joint modeling strategy based on BART leads to superior performance compared to other methods based on the cross-validated AUC. **B**. Metabolomics was the top contributor in the prediction of IBD status, as revealed by the estimated layer weights from the stacked layer. **C**. Top 20 metagenomics species (left) and metabolite compounds (right) based on variable importance scores in the integrative Human Microbiome Project (iHMP). Each row illustrates a per-feature variable inclusion proportion score representing the proportion of times each predictor is chosen as a splitting rule divided by the total number of splitting rules appearing in the model. In addition to reporting previously published differentially abundant bacterial species and metabolite compounds, *IntegratedLearner* is able to extract ‘a set of weakly predictive’ features that are together strongly predictive of IBD disease status giving rise to testable hypotheses warranting further investigation and offering actionable insights.

### 3.4 *IntegratedLearner* adapts to the missing data layers while maintaining accuracy in incomplete multi-omics data

We conclude our case studies by applying *IntegratedLearner* to another public multi-omics dataset ^5^ to showcase the utility of the method in situations when not all the layers from the training set are available in the validation set, a critical issue in the integrative analysis of multi-omics data. Missing layers can be caused by features failing to be measured in some of the samples, or by detection difficulties in a subset of samples. To this end, we considered longitudinal multi-omics data from pregnant women receiving routine antepartum care at the Lucile Packard Children’s Hospital in Stanford, CA, USA ^5^. The aim of this observational study was to determine a precise chronology of pregnancy-related metabolomic, proteomic, and immunologic adaptations in venous peripheral blood samples collected serially during the last 100 days of pregnancy from 112 participants leading up to 6286 combined features per sample (**Table 1**). The continuous outcome variables of this study included (i) the day of labor onset which is defined as the day of admission for spontaneous labor (contractions occurring at least every 5 min, lasting *>*1 min, and associated with cervical change), and (ii) gestational age which was determined by the best obstetrical estimate as recommended by the American College of Obstetricians and Gynecologists ^4^. Similar to Ghaemi *et al.* (2019) ^4^, we employed a one-subject-leave-out cross-validation to build prediction models in this dataset.

To address the missing data layer issue in the validation set, we adopted a two-stage approach in which we first learned the yer-wise weights based on all available layers in the training set using the *IntegratedLearner* default algorithm. In order to apply the trained model to the validation set with a missing proteomic layer, we simply re-learned the meta-learner to estimate the updated weights based on the matching layers in both training and validation cohorts. Note that, this step avoids re-training the individual base-learners which not only saves computing time but also optimally utilizes all available data points in both training and validation cohorts. This is in sharp contrast to existing methods which must only use the matching layers across training and validation sets to generate final predictions and remove any non-matching layers during training. As before, *IntegratedLearner* remains the best-performing method across learning tasks for this dataset, where it outperforms the LASSO stacked generalized approach used in the original study (**Figs. 5-6**), while also enabling credible interval estimation, layer-wise weight calculation, and feature importance score quantification, as before (**Figs. S5-S6**). Taken together, these findings thus confirm that *IntegratedLearner* is able to adapt to complex study designs and missing data layers, facilitating inference even with partially available data.

**FIGURE 5.**
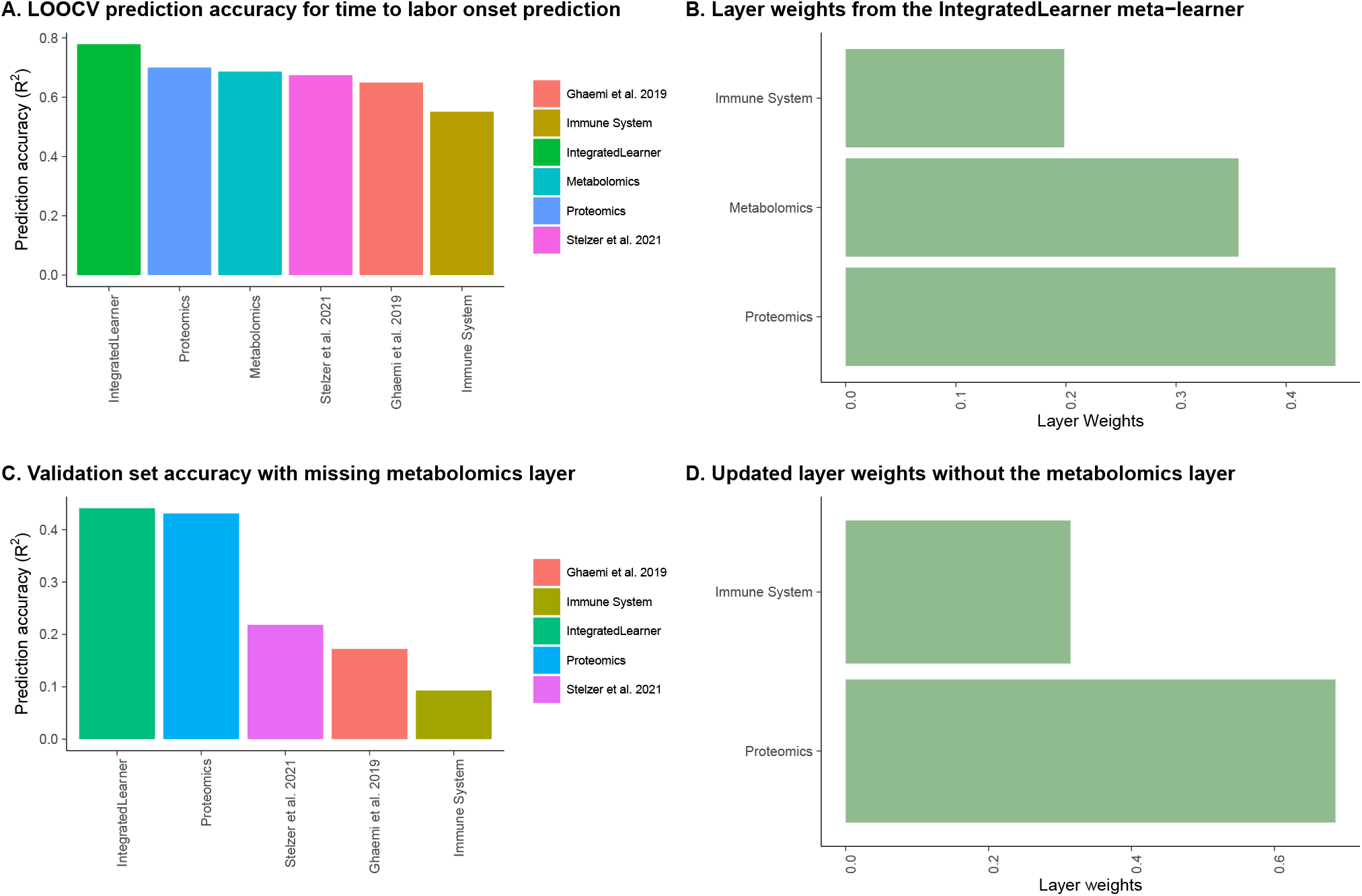
*IntegratedLearner* adapts to complex study designs with missing data layers in the validation set for time to labor onset prediction. A. We consider the problem of predicting the time to labor based on longitudinal multi-omics measurements with missing data layer in the validation set ^5^. In the training set, *IntegratedLearner* remains one of the bestperforming methods both across layers and combined, outperforming both Stelzer *et al.* (2021) ^5^ and Ghaemi *et al.* (2019) ^4^. **B**. Proteomics was the top contributor in the prediction of labor onset as revealed by the estimated *IntegratedLearner* layer weights. **C**. In the absence of a specific data layer in the validation set, *IntegratedLearner* re-learns the weights for the available layers without re-training the individual base-learners and achieves superior results compared to existing non-adaptive methods. **D**. Once again, proteomics remained the top contributor in the prediction of time to labor based on the updated weights.

**FIGURE 6.**
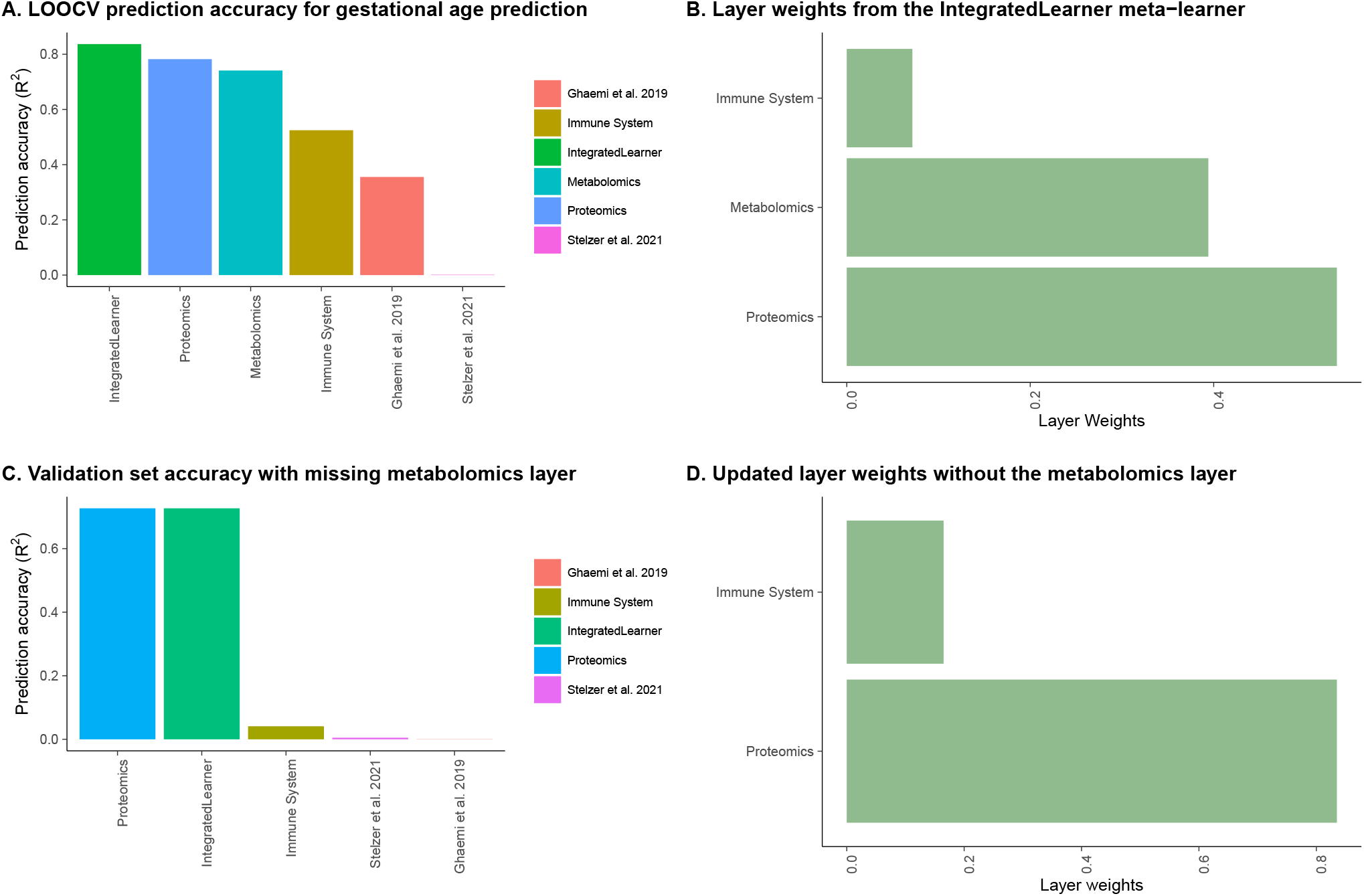
*IntegratedLearner* adapts to complex study designs with missing data layers in the validation set for gestational age prediction. A. We consider the problem of predicting the gestational age based on longitudinal multi-omics measurements with missing data layer in the validation set ^5^. In the training set, *IntegratedLearner* remains one of the best-performing methods both across layers and combined, outperforming both Stelzer *et al.* (2021) ^5^ and Ghaemi *et al.* (2019) ^4^. **B**. Proteomics was the top contributor in the prediction of gestational age as revealed by the estimated *IntegratedLearner* layer weights. **C**. In the absence of a specific data layer in the validation set, *IntegratedLearner* re-learns the weights for the available layers without re-training the individual base-learners and achieves superior results compared to existing non-adaptive methods. **D**. Once again, proteomics remained the top contributor in the prediction of gestational age based on the updated weights.

### 3.5 Simulation Studies

Beyond these real-world applications, the usefulness of *IntegratedLearner*, of course, depends on its accuracy in retrieving ground-truth signals from simulated multi-omics datasets. As noted by others ^19^, gold standard datasets that incorporate multiple omics and provide unbiased ground truth are a prerequisite for proper systematic evaluation of multi-omics methods and the best way to validate and study the properties of the proposed method will be to take real multi-omics data obtained from many individuals and simulate phenotypes based on these multi-omics features, making assumptions only about the data generation process. To characterize this effect, following an anonymous reviewer, we conducted a simulation study by mimicking real multi-omics data where we generate multiple interrelated data types with realistic intra- and inter-relationships based on the DNA methylation, mRNA gene expression, and protein expression from an ovarian cancer study in The Cancer Genome Atlas (TCGA) using the R package *InterSIM* ^20^. To be as realistic as possible, we used the simulated multi-omics features (i.e., the ‘s) from *InterSIM* to generate a continuous outcome, and performed a range of simulation studies by varying the data generation mechanisms incorporating diverse linear and non-linear effects, signal-to-noise ratios, and sample sizes (**Table 2**).

**TABLE 2.** Comparison of *IntegratedLearner* and published multi-omics methods in terms of out-of-sample prediction accuracy. Values are median out-of-sample *R*^2^ and standard deviations (in parentheses) summarized over 100 simulation runs. For each simulation scenario, the best method is boldfaced.

| Simulation | Sample Size | SNR | Franzosa *et al.*<sup>6</sup> | Ghaemi *et al.*<sup>4</sup> | Stelzer *et al.*<sup>5</sup> | *IntegratedLearner* |
| --- | --- | --- | --- | --- | --- | --- |
| Simulation 1<br>(Highly non-linear) | 200 | 1 | 0.210 (0.202) | 0.226 (0.306) | 0.234 (0.307) | **0.348 (0.308)** |
|  |  | 5 | 0.330 (0.202) | 0.383 (0.327) | 0.394 (0.329) | **0.556 (0.293)** |
|  |  | 10 | 0.367 (0.203) | 0.391 (0.334) | 0.419 (0.337) | **0.637 (0.291)** |
|  | 500 | 1 | 0.302 (0.212) | 0.304 (0.272) | 0.303 (0.273) | **0.409 (0.319)** |
|  |  | 5 | 0.471 (0.216) | 0.435 (0.315) | 0.442 (0.316) | **0.666 (0.303)** |
|  |  | 10 | 0.501 (0.207) | 0.448 (0.327) | 0.464 (0.328) | **0.752 (0.290)** |
|  | 1000 | 1 | 0.369 (0.241) | 0.337 (0.302) | 0.343 (0.303) | **0.516 (0.349)** |
|  |  | 5 | 0.538 (0.202) | 0.565 (0.312) | 0.568 (0.315) | **0.763 (0.291)** |
|  |  | 10 | 0.596 (0.180) | 0.634 (0.314) | 0.634 (0.318) | **0.806 (0.249)** |
| Simulation 2<br>(Highly linear) | 200 | 1 | 0.186 (0.185) | 0.285 (0.271) | 0.292 (0.270) | **0.350 (0.258)** |
|  |  | 5 | 0.288 (0.179) | 0.468 (0.265) | 0.467 (0.266) | **0.509 (0.225)** |
|  |  | 10 | 0.310 (0.175) | 0.502 (0.272) | 0.503 (0.266) | **0.528 (0.224)** |
|  | 500 | 1 | 0.307 (0.165) | **0.384 (0.216)** | 0.378 (0.217) | **0.384 (0.246)** |
|  |  | 5 | 0.431 (0.140) | 0.603 (0.222) | **0.605 (0.222)** | 0.572 (0.182) |
|  |  | 10 | 0.458 (0.138) | 0.650 (0.228) | **0.652 (0.228)** | 0.612 (0.175) |
|  | 1000 | 1 | 0.404 (0.160) | 0.449 (0.231) | 0.447 (0.231) | **0.494 (0.255)** |
|  |  | 5 | 0.531 (0.109) | 0.680 (0.228) | **0.683 (0.228)** | 0.651 (0.171) |
|  |  | 10 | 0.561 (0.102) | **0.723 (0.233)** | **0.723 (0.233)** | 0.678 (0.156) |

For each of these settings, we generate 100 replicates of training and test datasets, each with 658 features across three layers (131 gene expression, 367 methylation, and 160 protein features) and vary the signal-to-noise ratio (SNR) from low (1) to moderate (5) to high (10) for varying sample sizes (*n* = 200, *n* = 500, and *n* = 1000), where SNR is defined as 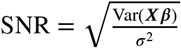.

We simulate data from the true model **y** = *f*_*0*_ (**X**) + ϵ, ϵ ∼ N(**0, σ**^2^ *I*)where σ^2^ is chosen such that the desired level of SNR is achieved. We consider the following data generation mechanisms:

1. **Simulation 1 (“Highly non-linear”)**: In this example, *f*_0_(**X**) = 10 sin(π*X*_1_*X*_2_) + 20(*X*_3_− 0.5)^2^ + 10*X*_4_+ 5*X*_5_, which includes non-linearity and interaction effects among five randomly chosen multi-omics features (*X*_1_, … , *X*_5_). The function *f*_0_, proposed initially by Friedman (1991) ^21^, has non-linear dependence on the first three variables *X*_1_, *X*_2_and *X*_3_, linear dependence on *X*_4_and *X*_5_, and incorporates a non-linear interaction between *X*_1_and *X*_2_.
2. **Simulation 2 (“Highly linear”)**: Similar to Simulation 1 except *f*_0_(**X**) = 10 sin(π*X*_1_*X*_2_) + 20(*X*_3_− 0.5)^2^ + 10*X*_4_+ 5*X*_5_+ **X *β*,** where the first five elements of *β* are zero and only 10% of the remaining coefficients are non-zero, drawn randomly from a log-normal distribution such that it leads to realistically varying log_2_-fold changes between (−3, 3) representing both modest (e.g., *<* 2-fold differences) and strong (e.g., 8-fold) effect sizes using the Splatter simulation framework ^22,23^.

The simulation results clearly demonstrate that *IntegratedLearner* continues to stand out as one of the most effective methods across a diverse range of simulation parameters, a phenomenon that was consistent across all simulation scenarios (**Table 2**). Particularly, when dealing with strong non-linear effects (Simulation 1), *IntegratedLearner* outperforms both concatenated random forest ^6^ and stacked generalization-based approaches ^5,4^. In the presence of prominent linear effects (Simulation 2), *IntegratedLearner* remains competitive, exhibiting superior performance in 5 out of 9 scenarios. It’s worth noting that the stacked generalization models consistently exhibit robust performance, outperforming concatenation-based methods across all scenarios, in agreement with previous findings ^5,4^. These results thus underscore the advantage of employing a Bayesian machine learning technique to combine strengths across multi-level omics data and the importance of incorporating uncertainty in the integration process.

The backbone of the *IntegratedLearner* methodology is the R package *bartMachine* ^24^, which is a fast implementation of BART, significantly faster than other BART implementations. As a result, the implementation of the *IntegratedLearner* workflow provides an on-par computation time compared to published frequentist machine learners, especially given that the approach couples MCMC and CV processes, making it an attractive alternative for a variety of practical applications (**Table 3**).

**TABLE 3.**
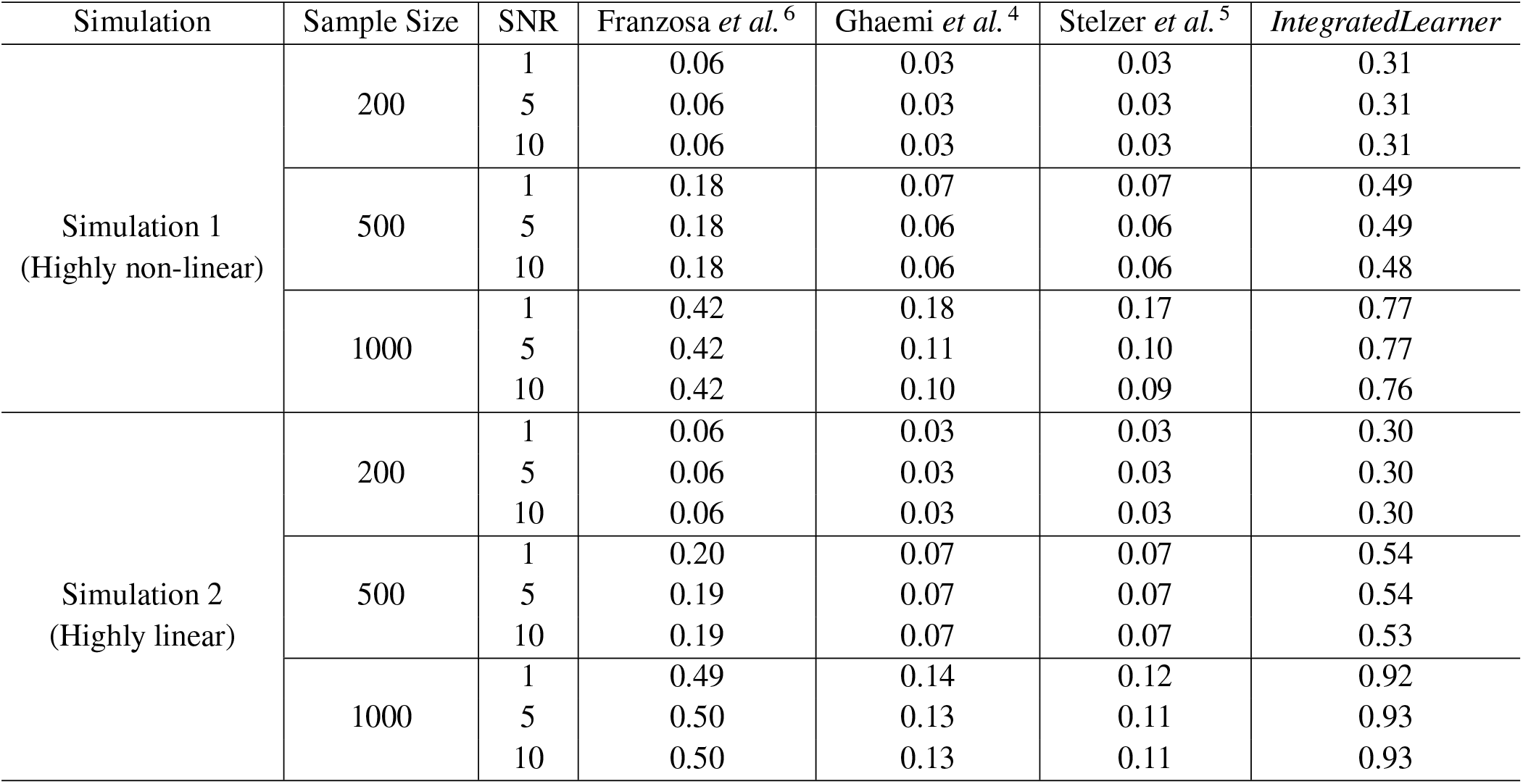
Comparison of average computing time (in minutes). Values are the median time summarized over 100 iterations.

| Simulation | Sample Size | SNR | Franzosa *et al.*<sup>6</sup> | Ghaemi *et al.*<sup>4</sup> | Stelzer *et al.*<sup>5</sup> | *IntegratedLearner* |
| --- | --- | --- | --- | --- | --- | --- |
| Simulation 1<br>(Highly non-linear) | 200 | 1 | 0.06 | 0.03 | 0.03 | 0.31 |
|  |  | 5 | 0.06 | 0.03 | 0.03 | 0.31 |
|  |  | 10 | 0.06 | 0.03 | 0.03 | 0.31 |
|  | 500 | 1 | 0.18 | 0.07 | 0.07 | 0.49 |
|  |  | 5 | 0.18 | 0.06 | 0.06 | 0.49 |
|  |  | 10 | 0.18 | 0.06 | 0.06 | 0.48 |
|  | 1000 | 1 | 0.42 | 0.18 | 0.17 | 0.77 |
|  |  | 5 | 0.42 | 0.11 | 0.10 | 0.77 |
|  |  | 10 | 0.42 | 0.10 | 0.09 | 0.76 |
| Simulation 2<br>(Highly linear) | 200 | 1 | 0.06 | 0.03 | 0.03 | 0.30 |
|  |  | 5 | 0.06 | 0.03 | 0.03 | 0.30 |
|  |  | 10 | 0.06 | 0.03 | 0.03 | 0.30 |
|  | 500 | 1 | 0.20 | 0.07 | 0.07 | 0.54 |
|  |  | 5 | 0.19 | 0.07 | 0.07 | 0.54 |
|  |  | 10 | 0.19 | 0.07 | 0.07 | 0.53 |
|  | 1000 | 1 | 0.49 | 0.14 | 0.12 | 0.92 |
|  |  | 5 | 0.50 | 0.13 | 0.11 | 0.93 |
|  |  | 10 | 0.50 | 0.13 | 0.11 | 0.93 |

## 4 DISCUSSION

We have developed and validated an integrated Bayesian machine learning method for multi-omics prediction and classification. The resulting framework enables the prediction of continuous and discrete binary outcomes from both cross-sectional and longitudinal multi-omics biomarkers with or without covariates. Philosophically, *IntegratedLearner* belongs to the class of several recently proposed integrative models that leverage multiple omics layers or datasets, providing a more rigorous approach to correcting for single-omics biases than earlier methods ^6,3,4^. Unlike existing frequentist methods that summarize predictions using a single point estimate, *IntegratedLearner* yields uncertainty estimates (i.e., credible intervals) of the prediction and model parameters in addition to reporting a small set of interpretable features for follow-up experiments. Numerical results show that *IntegratedLearner* outperforms published methods in estimation and prediction while confirming established results and suggesting novel ones for future validation as showcased in four public multi-omics real data applications. *IntegratedLearner* is open source, modular, and convenient to be extended to other multi-table input data without compromising on either theoretical guarantee or empirical performance ^25^, making it a promising tool for more accurate and personalized clinical decision-making from multiple datasets or studies.

Joint modeling using a two-stage approach provided one of the first use cases of incorporating longitudinal biomarkers in existing integrated machine learning frameworks. In particular, the two-stage approach allowed us to capture the dynamic effects of the time-varying biomarkers in the form of covariate-adjusted random effects, not readily achievable by existing cross-sectional frameworks. Further, the residualization strategy embedded in our framework enables zeroing in on the mechanistic drivers among the detected predictive features by minimizing the impact of residual confounding. Specifically, it highlights putative mechanistic features that are either (i) mutually predictive of the outcome or (ii) covary independently of their mutual covariation with the outcome, or (iii) both. Likewise, the identified features from the above-mentioned residualized prediction model can be interpreted as ‘potentially causal’ decorrelated features worthy of further investigation, facilitating a better downstream characterization of the identified biological entities.

To the best of our knowledge, this is one of the first works utilizing the BART for multi-omics data integration, which, we strongly believe, will open up new avenues for future methodological advances. For example, the current implementation of *IntegratedLearner* only allows for the missingness of entire data layers. However, for situations with missing measurements for a subset of variables within given layers, *IntegratedLearner* can be naturally extended to accommodate such missingness ^26^.

To assess the relative significance of predictors across various modalities, a potential approach involves merging the inferred layer weights from the meta-learner with the variable importance scores obtained from the single-layer BART models. Let’s denote the estimated layer weights for the *k*^*th*^ layer as 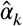, *k* = 1, … , *K*. Additionally, let the variable importance score for variable *j* in layer *k* be denoted by 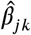, *j*= 1, … , *p*_*k*_, *k* = 1, … , *K*, where *p*_*k*_ denotes the number of predictors in *k*^*th*^ layer. Then, we can calculate the relative importance of a predictor *x*_*jk*_ in the `IntegratedLearner` model by 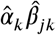, which quantifies the relative importance of variables from different modalities. Visual representation of the weighted feature importance scores, as defined above, is provided in **Figs. S7–S11**, showcasing the application of these scores to the case studies outlined in this manuscript.

As a cautionary note, we consider this as only a very preliminary evaluation of the potential usefulness of the weighted feature importance scores. Despite our encouraging findings, further research is definitely needed. Similarly, within-layer variable selection can be attained by enforcing sparsity within each individual layer ^27,28,29^. It’s worth acknowledging that incorporating prior information regarding the importance of specific data layers and/or variables within those layers, akin to certain existing frequentist methods ^3^, presents a challenge that remains unsolved and open for exploration in the upcoming years.

Limitations of the current method include, first, its restriction to building base-learners on a per-layer basis. As a result, the model currently does not share information between layers during the first-stage learning, as would be the case in a joint model, albeit at the expense of biased inference and extra computing time ^4^. We, therefore, hope that future work could further finetune the model by incorporating layer-wise correlations during the first-stage learning, building upon similar developments in other disciplines such as spatial statistics ^30^. Another extension opportunity entails considering an ensemble learner for each individual layer to further improve prediction although at the expense of interpretability ^12^ or use the capability of conformal prediction ^31,32^ and/or explainable AI ^33^ to further improve uncertainty quantification. Direct incorporation of random effects for repeated measures and clustered observations offers another promising avenue to integrate longitudinal biomarkers ^34,35^. Finally, and relatedly, it is not straightforward to incorporate any type of prior immunological knowledge (e.g., regulatory networks) into the Bayesian paradigm. Recently, Culos *et al.* (2020) ^9^ considered a Bayesian elastic net prior incorporating prior knowledge by means of a carefully constructed tensor of biologically consistent immunological features. A similar strategy could be considered for the *IntegratedLearner* base-learners during prior elicitation, wherein, a literature-curated tensor can be used to further refine the training and validation process. Although we have only considered the setup of continuous and binary outcomes, the extension to count or survival outcome prediction and multi-label classification is immediate ^36,37,38^.

It is to be noted that our per-feature BART model captures interactions within the same modality but does not explicitly model interactions between features across modalities. To address this limitation, we need to create pairwise or higher-order interaction matrices between features across modalities and feed them into *IntegratedLearner*. To address the computational bottleneck associated with high-dimensional interactions ^39^, one can identify latent factors that explain variation within or across multi-omics datasets and use them as input for subsequent model building ^40,41^. Additional research is needed to determine the generalizability of this approach in various diversified scenarios which we consider as a plausible future direction. In a similar vein, one of the use cases of our method entails the prediction of a cross-sectional outcome based on longitudinal multiomics measurements, where the goal of the analysis is to provide, for each subject, an estimated classification probability or prediction ^14^. Although the two-stage modeling strategy described in Section 2.3 accounts for the repeated-measurement design, it does not explicitly model time. In order to explicitly model the longitudinal trajectories of the multi-omics biomarkers, one approach could be to include time as a predictor variable in a mixed-effects per-layer BART model ^34^ that allows the covariance structure to vary over time, similar in spirit to published mixed-effects random forest methods ^42,35^.

A critical issue in the integrative analysis of multi-omics data is the problem of missing values at both sample and feature levels, for example, as highlighted by multiple case studies considered here ^16,5^. While most current methods including *IntegratedLearner* are designed to be trained on all available data points across samples and layers, future methods must account for the increasing commonality of unequal sample sizes across omics layers, and more critically, heterogeneous signal-to-noise ratios across data types ^43^. Another consequential experimental design consideration for future multi-omics data integration tools is the widespread availability of hierarchical observational units across data tables (e.g., digital pathology ^44^, single-cell multiomics ^45^), wherein the single-omics modalities can be summarized at either individual- or instance-level (e.g., cells nested within individuals ^46^ or spots ^47^, as would be expected in a single-cell or a spatial transcriptomics and/or digital pathology study). Growing commonality of spatial omics and digital pathology data further means that existing tools must carefully refine the quality control of individual data types to enable a general mapping between pixel coordinates and molecular profiles, enabling analysts to relate image-level observations to omics measurements in a unified, platform-agnostic manner ^48,49,50^. Combined, such extensions will allow researchers to further improve multi-omics prediction and classification beyond homogeneous observational units, moving towards a more actionable blueprint for novel biomarker discovery in clinical and pre-clinical studies. We believe that the improved prediction, actionable feature selection, and accurate uncertainty quantification provided by *IntegratedLearner* represents an important step in this direction that can ultimately aid in the discovery of multi-omics-based therapeutic targets.

## CODE AVAILABILITY

The implementation of *IntegratedLearner* is publicly available with source code, documentation, tutorial, and as an R/Bioconductor package at https://github.com/himelmallick/IntegratedLearner. Analysis scripts for synthetic benchmarking and real data analyses are available from the first author upon request.

## DATA AVAILABILITY

Previously published data used in this study are appropriately cited in the main text as well as in the section. The detailed data summary is provided in **Table 1**. All processed data for the four case studies are available at https://github.com/himelmallick/IntegratedLearner.

## ACKNOWLEDGEMENTS

The authors extend their heartfelt gratitude to the dedicated editorial team and the anonymous referees for their insightful comments and suggestions. The authors would also like to express their deep appreciation to Drs. Andy Liaw (Merck Research Laboratories), Jia Kang (Merck Research Laboratories), and Christopher Woelk (Verge Genomics), whose invaluable feedback and insights have played a pivotal role in the refinement and enrichment of the *IntegratedLearner* methodology.

## APPENDIX

**TABLE S1.** AUC scores for various integrative classification methods for PRISM and iHMP datasets for classifying IBD status. For DIABLO, we used the default implementation (as available in the *mixOmics* R package) with an interaction coefficient of 0.1, as suggested by the authors ^3^.

| Method | PRISM |  | iHMP |
| --- | --- | --- | --- |
|  | Discovery | Validation |  |
| *IntegratedLearner* | 0.95 | **0.96** | **0.82** |
| Ghaemi *et al.* | 0.41 | 0.5 | 0.79 |
| Fransoza *et al.* | **0.96** | 0.89 | 0.78 |
| DIABLO | 0.76 | 0.71 | 0.77 |

**FIGURE S1.**
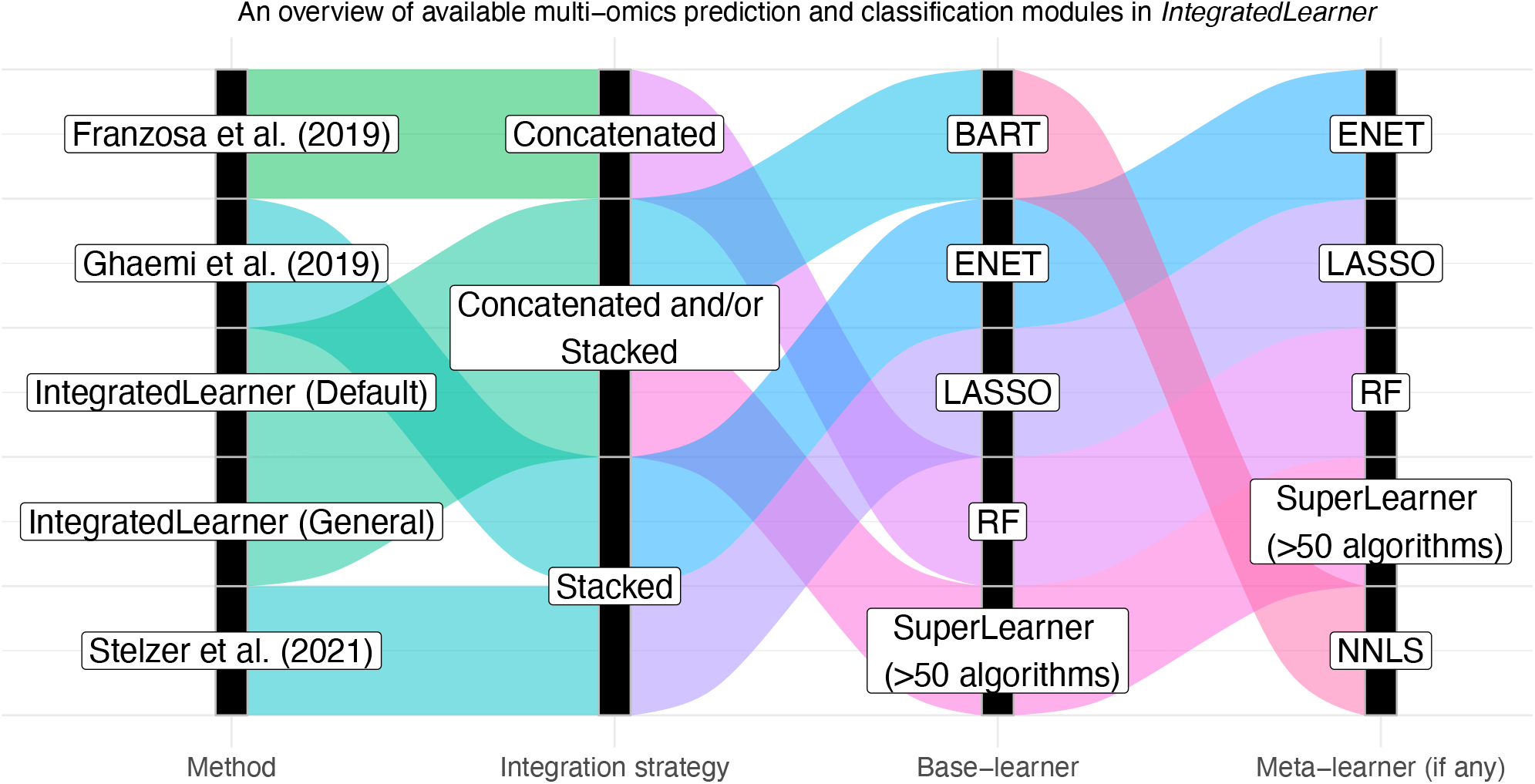
An overview of various integrative ML algorithms evaluated in the study. Most of the existing methods are directly implemented in *IntegratedLearner*. Unlike existing methods, *IntegratedLearner* is not married to a specific integration strategy or a machine learning algorithm. Instead, *IntegratedLearner* simplifies the multi-omics classification and prediction workflow by allowing an expanded combination of integration strategies and machine learning algorithms (>50 methods supported) based on the *SuperLearner* R package ^12^).

**FIGURE S2.**
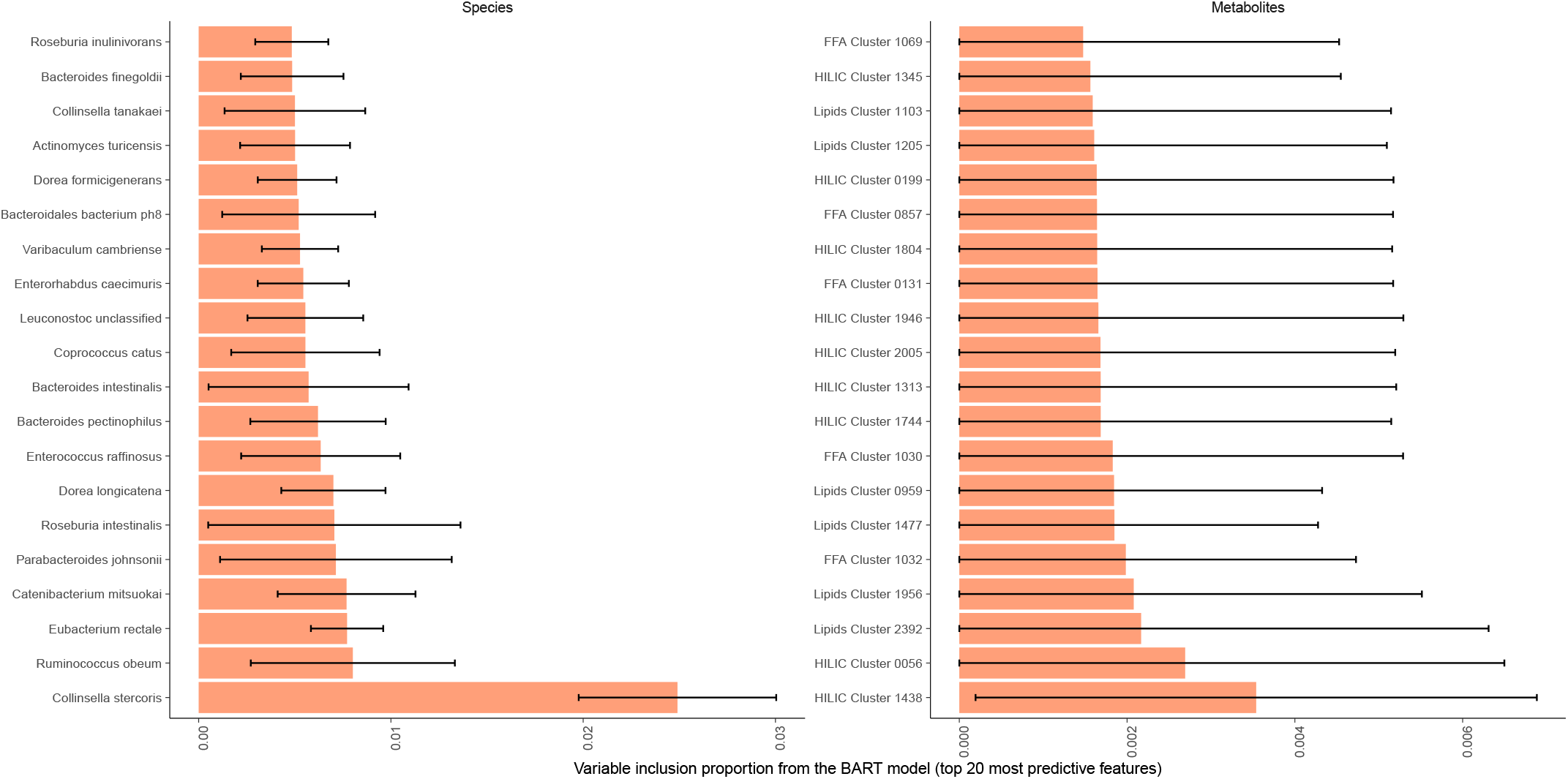
Feature importance scores of top 20 variables selected by the BART base-learners in the PRISM dataset ^6^. None of the 20 top metabolite features were annotated meaning that the identities of these metabolites are yet to be confirmed against laboratory standards.

**FIGURE S3.**
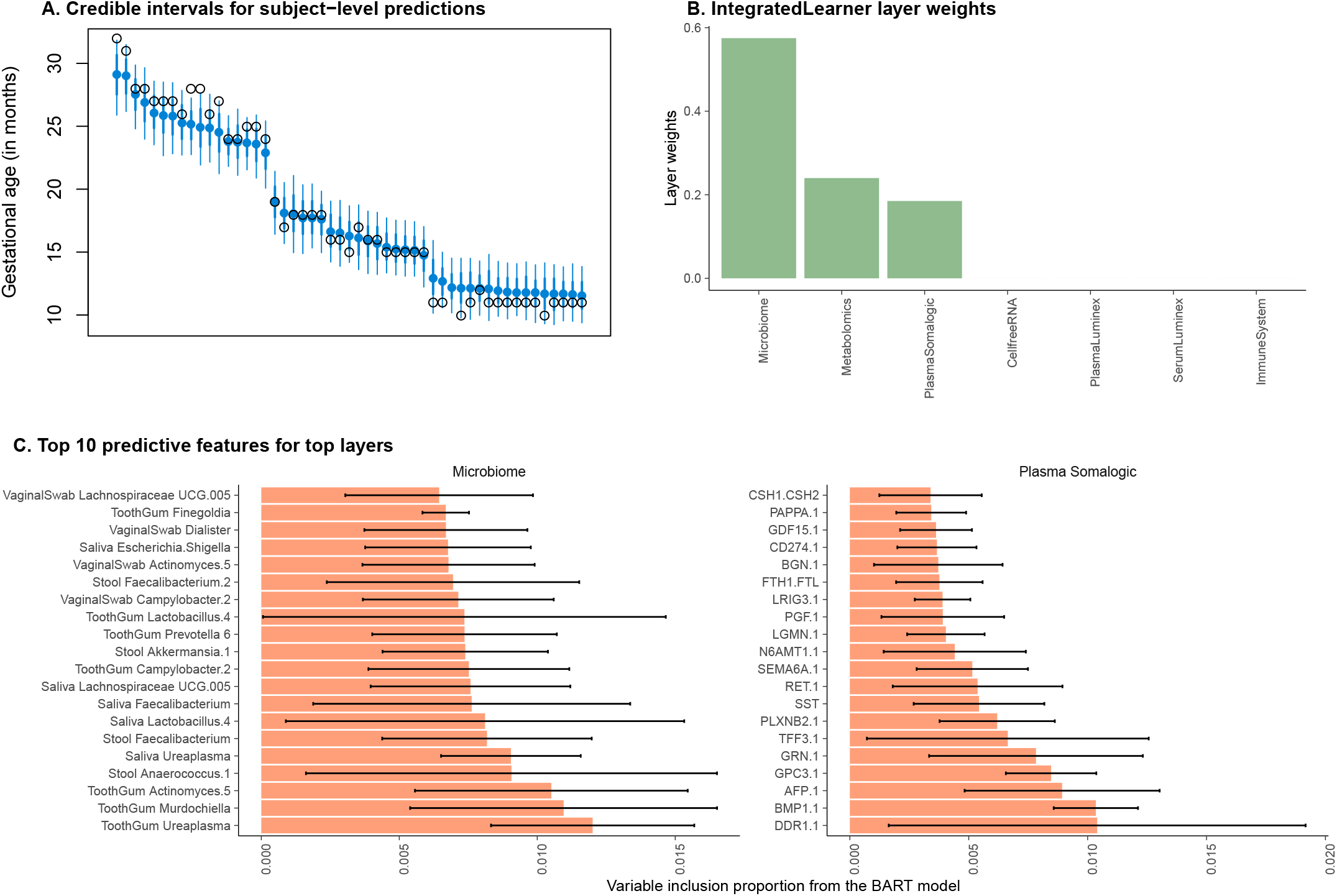
A. Our method produces tight credible intervals for gestational age in the pregnancy dataset ^4^. The thick lines represent 68% and the thin lines represent 95% credible intervals; the heights of ‘o’ represent the true labels and the heights of filled ‘o’ denote the predicted posterior median gestational age. **B**. Microbiome, metabolomics, and plasma SomaLogic were the top three contributors in the *IntegratedLearner*’s second-stage meta-learner as revealed by the estimated weights from the stacked layer. **C**. *IntegratedLearner* offers readily available feature importance scores as a byproduct of using the BART models as base-learners. We show the top 10 features selected by our model in two out of the top three layers for illustrative purposes.

**FIGURE S4.**
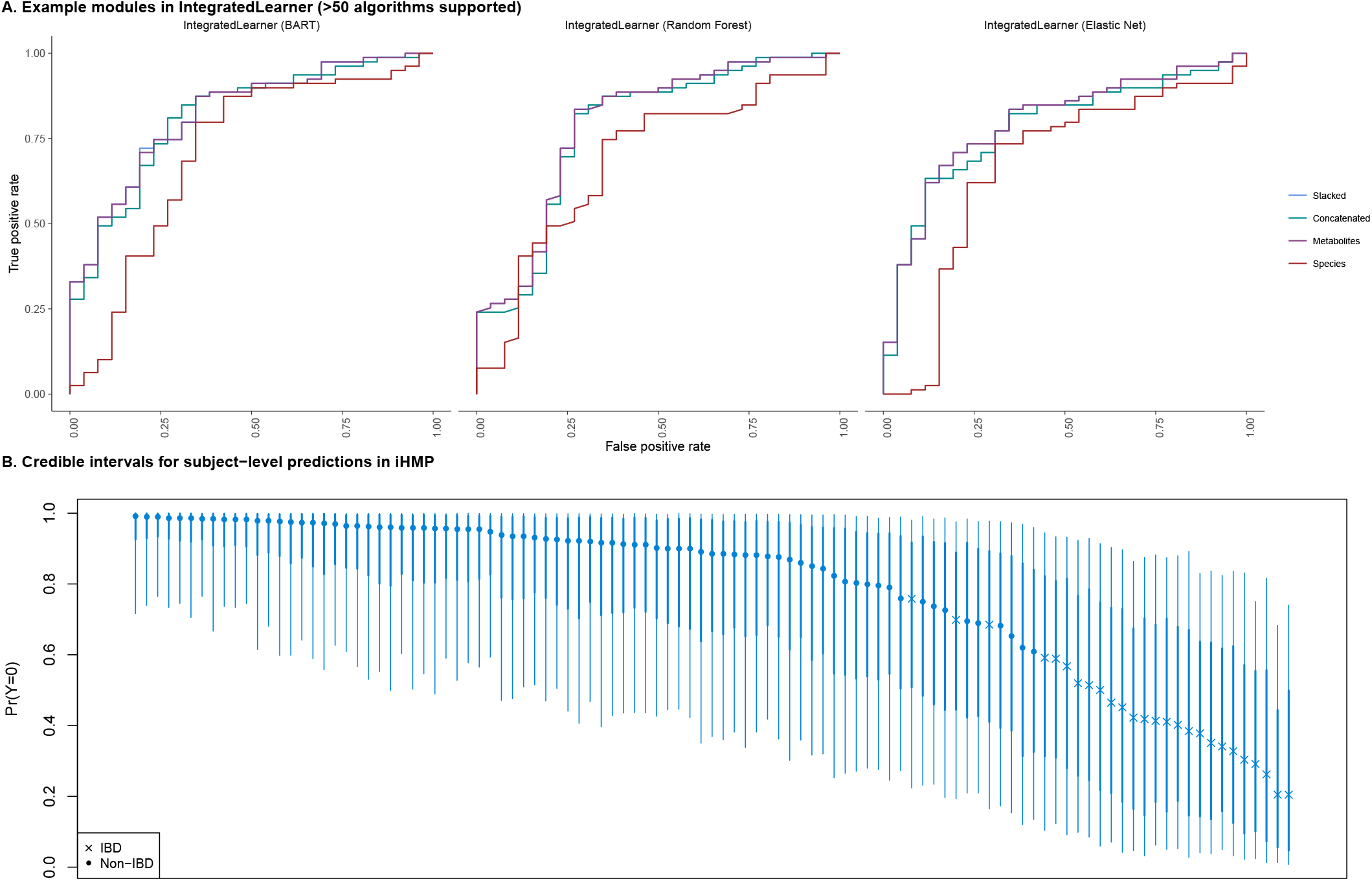
A. An illustration of *IntegratedLearner*’s two-stage longitudinal multi-omics classification modules for iHMP dataset ^16^. Apart from the default BART learners, a variety of other modules are available for the users and the software supports several other models to customize analysis for a specific study. **B**. *IntegratedLearner* automatically provides the posterior credible intervals. The thick lines represent 68% and the thin lines represent 95% credible intervals; ‘x’ or ‘o’ represent the true labels and their heights denote the predicted posterior median probabilities of no disease.

**FIGURE S5.**
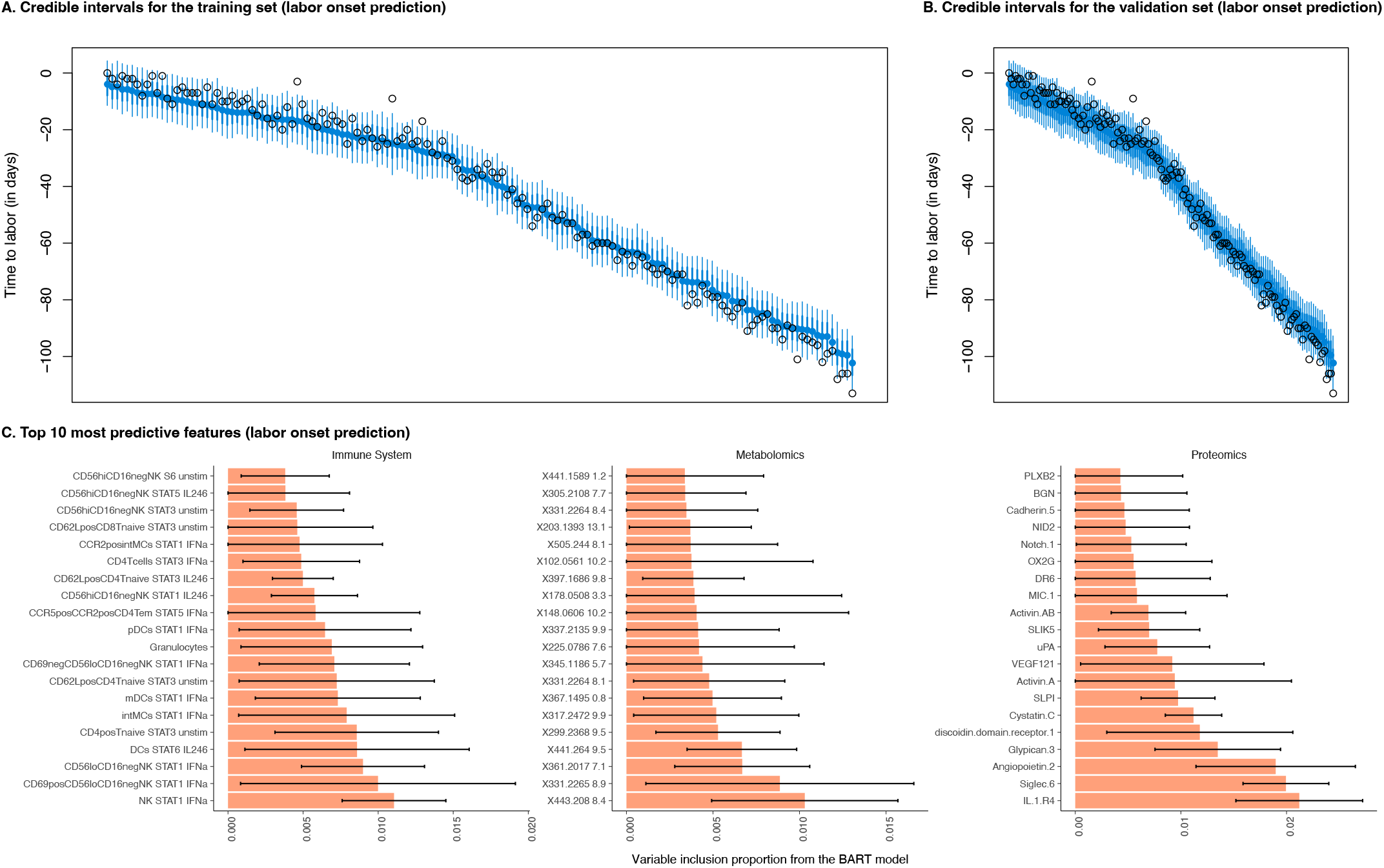
A and **B** present posterior credible intervals available for the training and validation set in the Stelzer *et al.* (2021) ^5^ dataset. **C** shows the top 10 predictive features of each of the three modalities in the Stelzer *et al.* (2021) ^5^ dataset when the outcome variable is time to labor onset.

**FIGURE S6.**
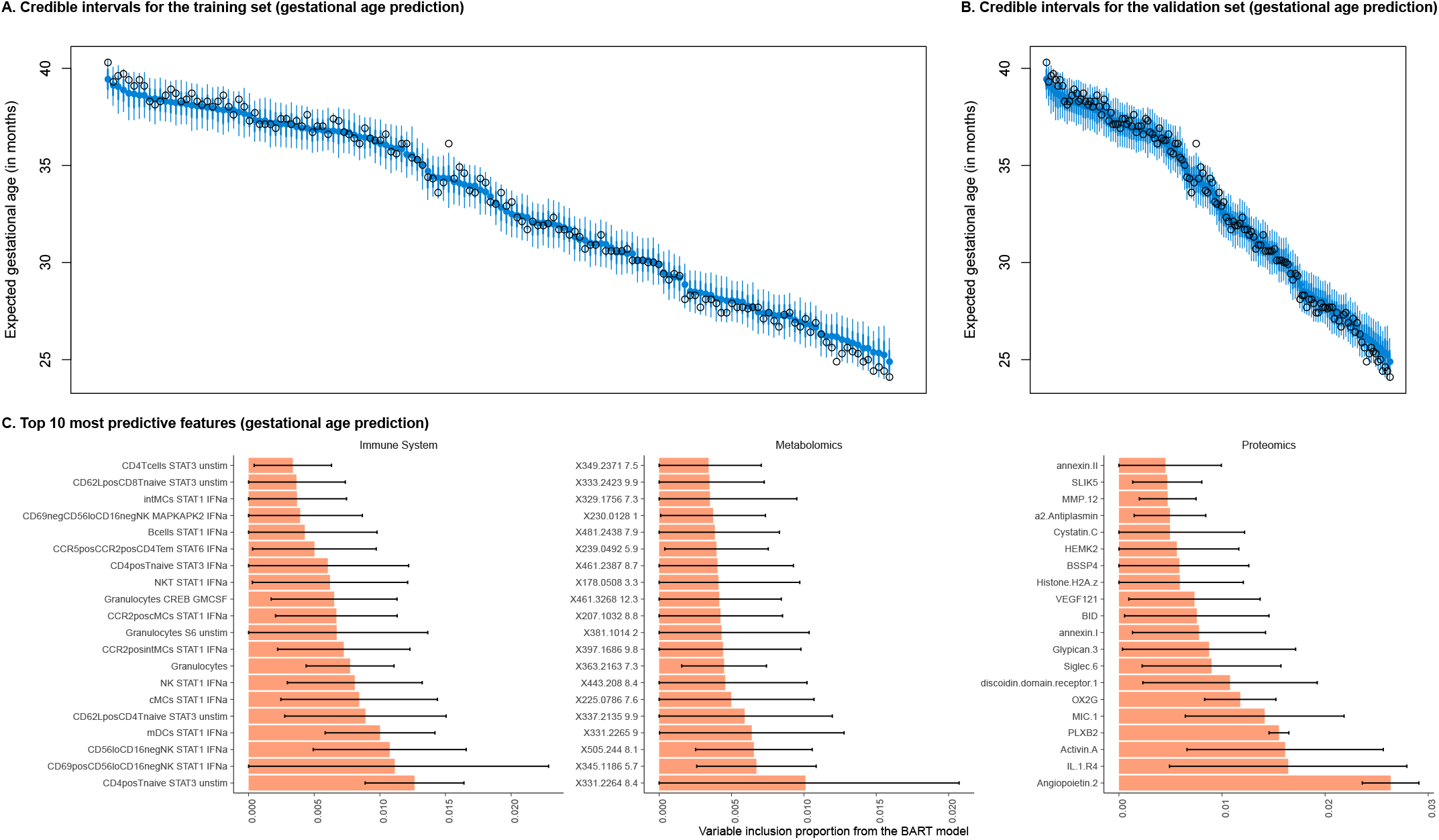
A and **B** present posterior credible intervals available for the training and validation set in the Stelzer *et al.* (2021) ^5^ dataset. **C** shows the top 10 predictive features of each of the three modalities in the Stelzer *et al.* (2021) ^5^ dataset when the outcome variable is expected gestational age for pregnant women in the study.

**FIGURE S7.**
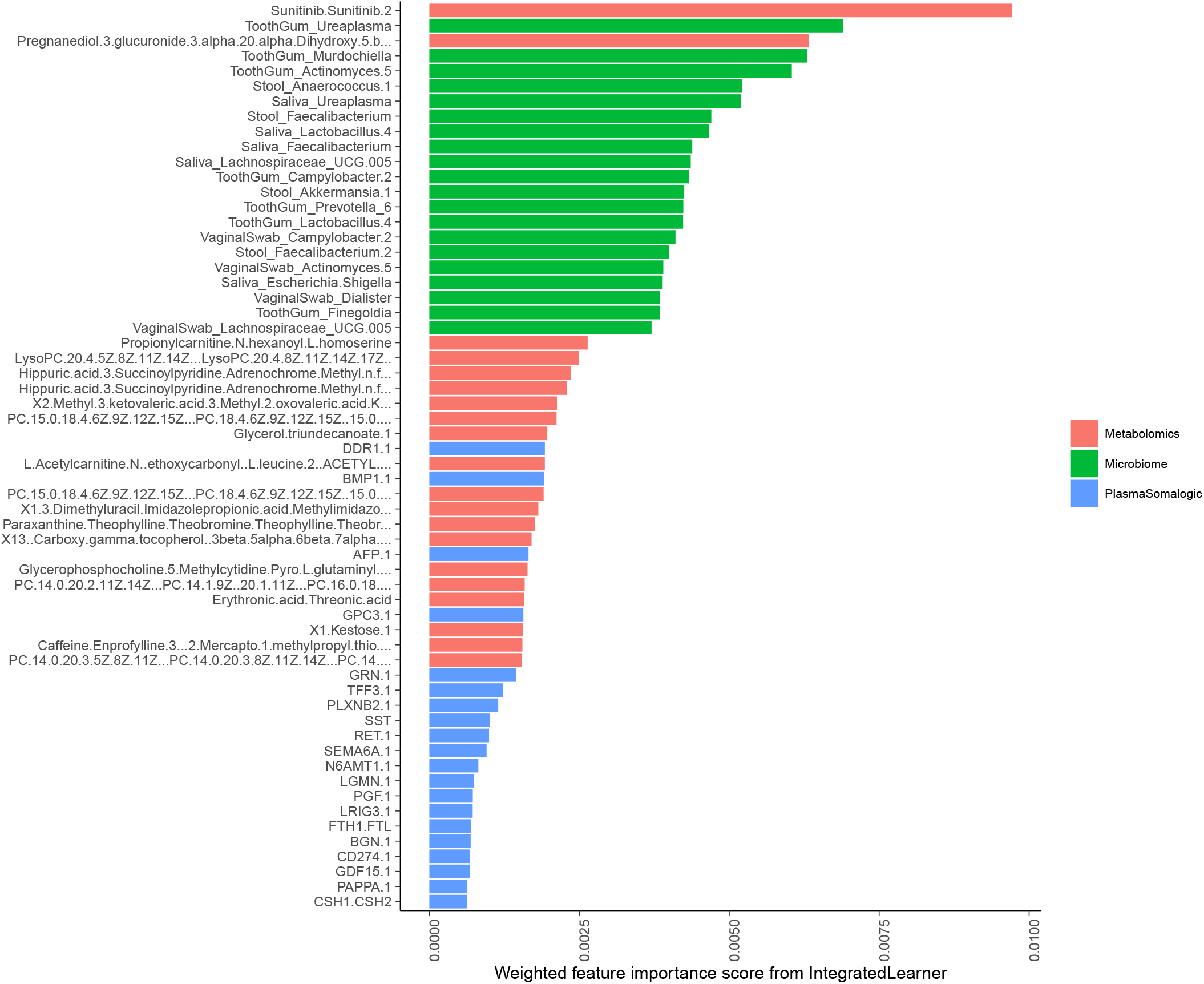
Weighted feature importance scores from the *IntegratedLearner* model for the pregnancy dataset ^4^. For visualization purposes, long feature names are truncated.

**FIGURE S8.**
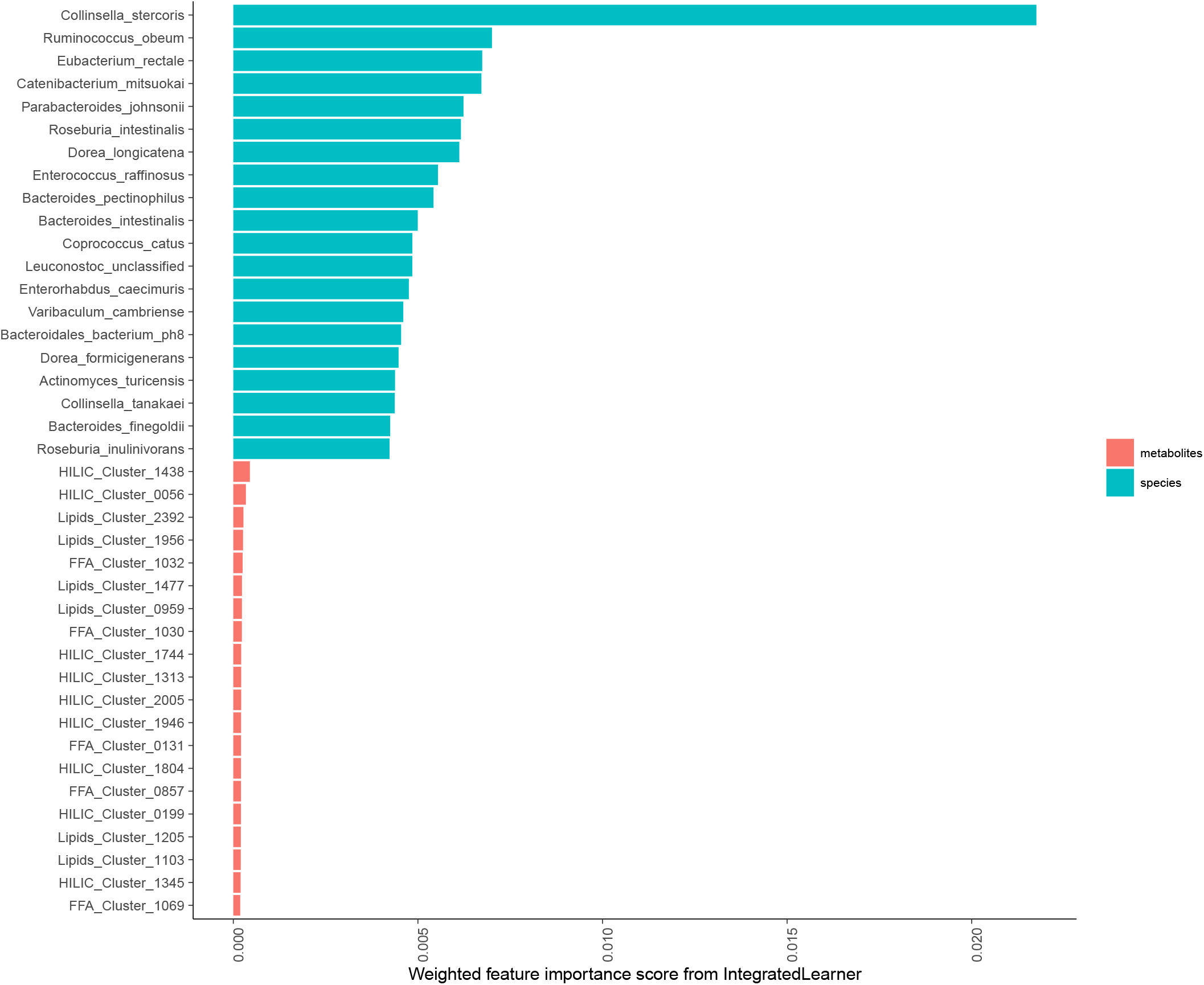
Weighted feature importance scores from the IntegratedLearner model for the PRISM dataset ^6^. For visualization purposes, long feature names are truncated.

**FIGURE S9.**
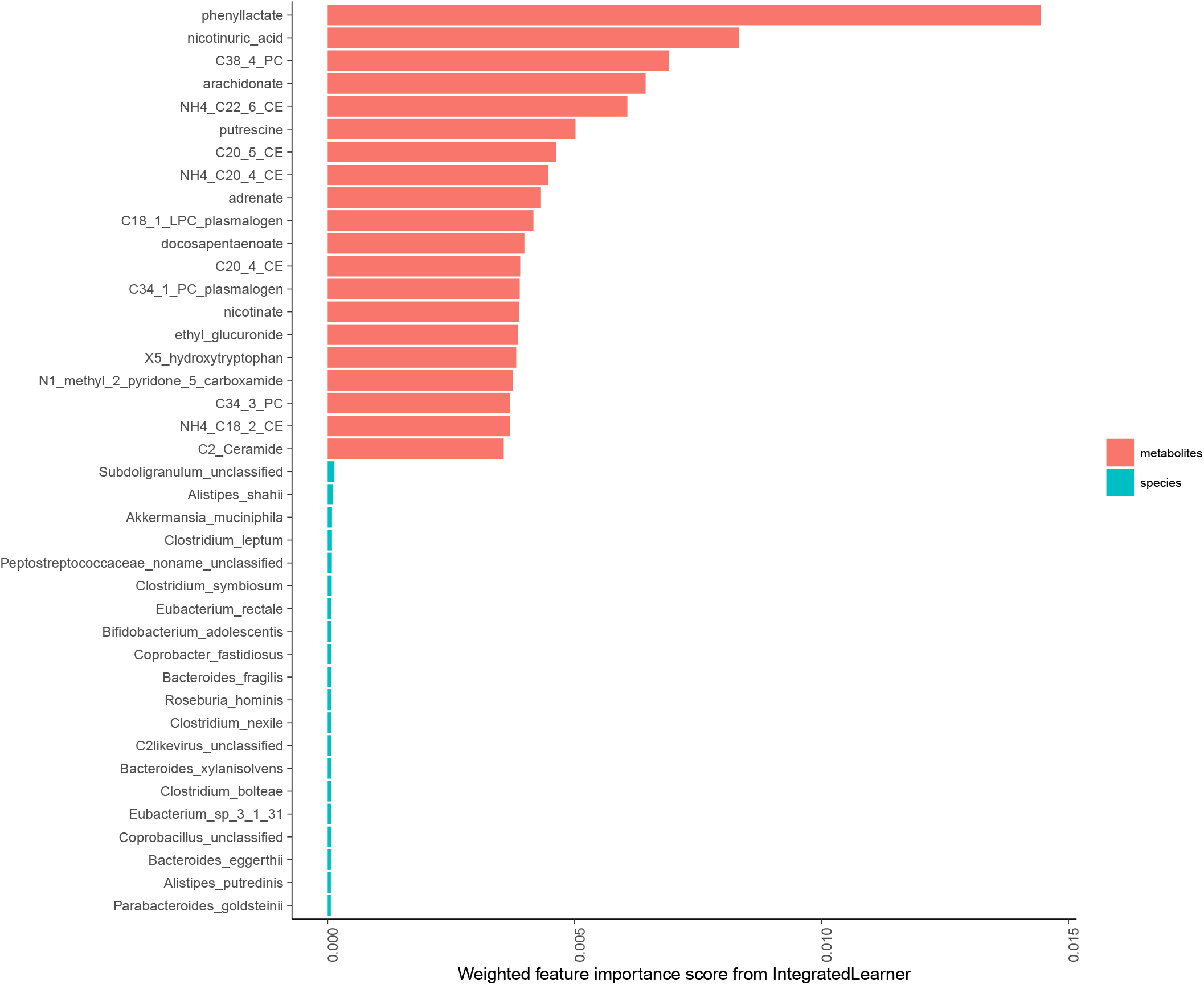
Weighted feature importance scores from the *IntegratedLearner* model for the iHMP dataset ^16^. For visualization purposes, long feature names are truncated.

**FIGURE S10.**
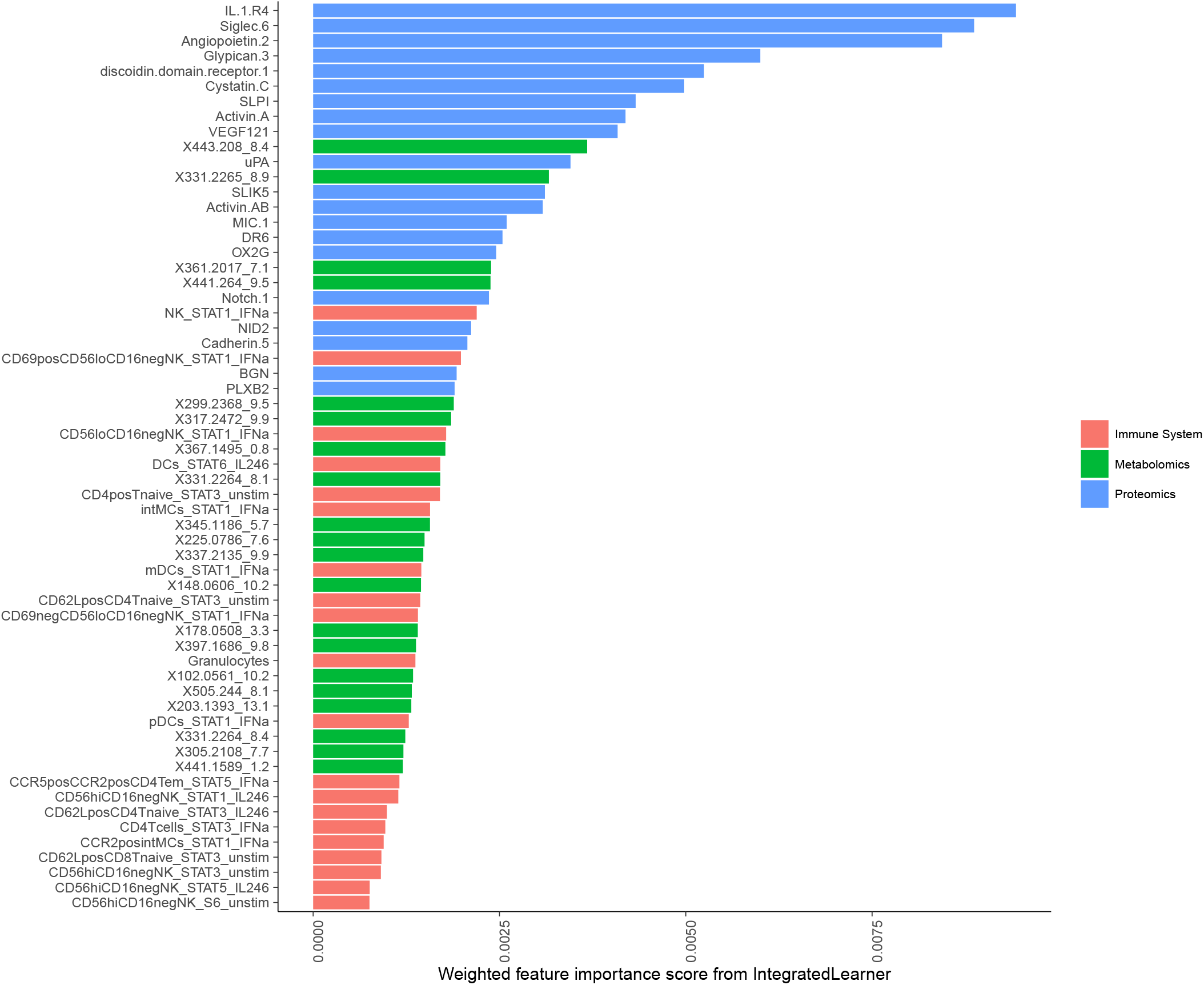
Weighted feature importance scores from the *IntegratedLearner* model for the Stelzer *et al.* (2021) dataset ^5^ (labor onset prediction). For visualization purposes, long feature names are truncated.

**FIGURE S11.**
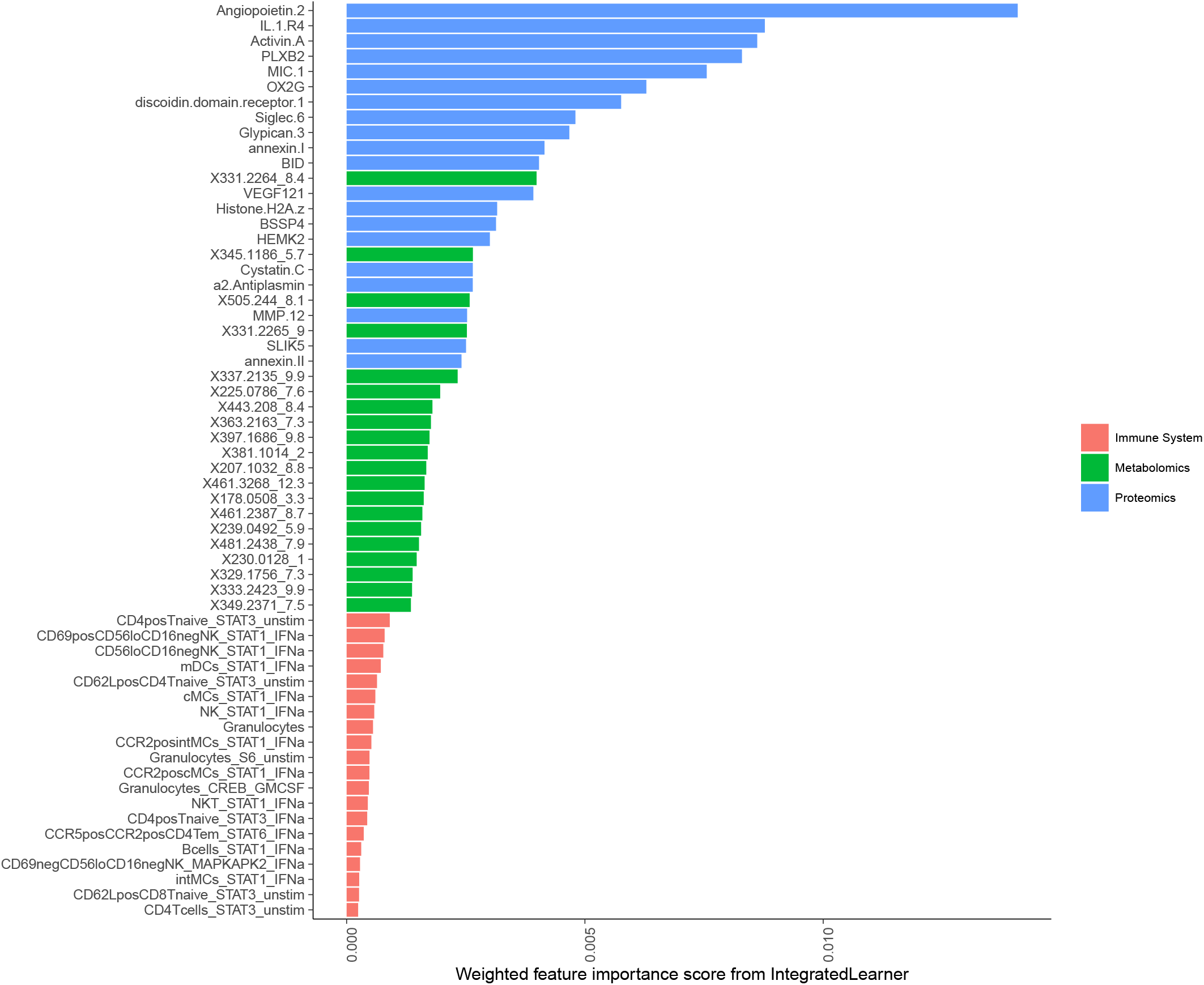
Weighted feature importance scores from the *IntegratedLearner* model for the Stelzer *et al.* (2021) dataset ^5^ (gestational age prediction). For visualization purposes, long feature names are truncated.

